# A kinetics-based model of hematopoiesis reveals extrinsic regulation of skewed lineage output from stem cells

**DOI:** 10.1101/2025.02.04.636388

**Authors:** Esther Rodríguez-Correa, Florian Grünschläger, Tamar Nizharadze, Natasha Anstee, Jude Al-Sabah, Vojtech Kumpost, Anastasia Sedlmeier, Congxin Li, Melanie Ball, Foteini Fotopoulou, Jeyan Jayarajan, Ian Ghezzi, Julia Knoch, Megan Druce, Theo Aurich, Marleen Büchler-Schäff, Susanne Lux, Pablo Hernández-Malmierca, Julius Gräsel, Dominik Vonficht, Anna Mathioudaki, Judith Zaugg, Ralf Mikut, Andreas Trumpp, Thomas Höfer, Daniel Hübschmann, Simon Haas, Michael D. Milsom

**Affiliations:** Division of Experimental Hematology, German Cancer Research Center (DKFZ), Heidelberg, Germany; Heidelberg Institute for Stem Cell Technology and Experimental Medicine (HI-STEM gGmbH), Heidelberg, Germany; Faculty of Biosciences, Heidelberg University, Heidelberg, Germany; Division of Stem Cells and Cancer, German Cancer Research Center (DKFZ), Heidelberg, Germany; Division of Theoretical Systems Biology, German Cancer Research Center (DKFZ), Heidelberg, Germany; Institute for Automation and Applied (IAI), Informatics, Karlsruhe Institute of Technology (KIT), Karlsruhe, Germany; Computational Oncology, Molecular Precision Oncology Program, National Center for Tumor Diseases (NCT) Heidelberg and German Cancer Research Center (DKFZ), Heidelberg, Germany; Molecular Systems Biology Unit, European Molecular Biology Laboratory, Heidelberg, Germany; Division of AI in Oncology, German Cancer Research Centre (DKFZ), Heidelberg, Germany; Department of Biomedicine, University Hospital Basel, University of Basel, Basel, Switzerland; DKFZ–ZMBH Alliance, Heidelberg, Germany; German Cancer Consortium (DKTK), Heidelberg, Germany; Innovation and Service Unit for Bioinformatics and Precision Medicine, German Cancer Research Center (DKFZ), Heidelberg, Germany; Division of Translational Precision Medicine, Institute of Human Genetics, Heidelberg University, Heidelberg, Germany; Berlin Institute of Health (BIH) at Charité Universitätsmedizin, Berlin, Germany; Berlin Institute for Medical Systems Biology, Max Delbrück Center for Molecular Medicine in the Helmholtz Association, Berlin, Germany; Precision Healthcare University Research Institute, Queen Mary University of London, London, UK

## Abstract

Residing at the top of the hematopoietic hierarchy, long-term hematopoietic stem cells (HSCs) are capable of self-renewal and sustained blood cell regeneration. Over the past decades, single-cell and clonal analyses have revealed substantial functional and molecular heterogeneity within this compartment, challenging the notion that self-renewal is inherently tied to balanced, multi-lineage blood production. However, a cohesive model that explains the relationships among these diverse HSC states remains elusive. Here, we combined single-cell transplantations of over 1,000 highly purified murine long-term HSCs with in-depth phenotyping of their clonal progeny to achieve a detailed, time-resolved understanding of heterogeneous reconstitution outcomes. We identified reconstitution kinetics as an overall unifying metric of HSC functional potency, with the most potent HSCs displaying the greatest delay in hematopoietic reconstitution. Importantly, a progressive acceleration in reconstitution kinetics was also associated with a gradual shift in mature cell production from platelet and erythro-myeloid bias to balanced, and eventually lymphoid bias. Serial single-cell transplantations of HSCs revealed a unidirectional acceleration in reconstitution kinetics accompanied by a gradual decline in functional potency of daughter HSCs, aligning diverse phenotypes along a linear hierarchical trajectory. Mathematical modeling, together with experimental modulation of lineage-biased blood production, demonstrated that apparent lineage biases actually arise from cell-extrinsic feedback regulation and clonal competition between slow- and fast-engrafting clones to occupy the limited compartment sizes of mature lineages. Our study reconciles multiple layers of HSC heterogeneity into a unifying framework, prompting a reevaluation of the meaning of lineage biases in both normal and diseased hematopoiesis, with broad implications for other regenerating tissues during development, homeostasis, and repair.

## Introduction

Hematopoietic stem cells (HSC) have canonically been perceived as a uniform entity with consistent self-renewal and multipotent characteristics^1^. However, over the course of the last two decades, numerous studies have characterized an ever-increasing spectrum of functional and molecular heterogeneity within this compartment^2,3^. Transplantation studies of variable immunophenotypically-defined populations demonstrated profound differences in their capacity to sustain hematopoiesis over time, identifying HSCs with long- or short-term repopulation capacity, as well as multipotent progenitors (MPPs) with transient engraftment and limited self-renewal^4^. Single-cell transplantation and barcoding experiments have shown that, even within the immunophenotypically homogenous long-term (LT)-HSC compartment, the quantitative and qualitative output of individual HSCs is highly heterogeneous^5–8^. In this context, multiple studies have reported pronounced biases of HSCs regarding the generation of distinct lineages of the hematopoietic system, including myeloid and lymphoid biased output, as well as individual HSCs that are capable of generating a more balanced multilineage reconstitution pattern^6,7,9,10^. More recent studies also identified platelet-biased HSCs, which appear to reside at the apex of the hematopoietic hierarchy^8,11–13^. While several physiological roles of lineage biases have been suggested^12,14^, the underlying mechanisms via which such biased blood production programs are established remain unknown. In line with the observed functional heterogeneity within the HSC pool, single-cell multi-omic profiling has also revealed significant molecular heterogeneity of HSCs, correlating with distinct stemness and lineage bias patterns^8,13,15–17^. These molecular analyses suggest a spectrum of heterogeneity along continuous gradients^17–20^. Together, these functional and molecular studies have challenged the classical model of hematopoiesis, which assumes HSCs are multipotent and homogeneous in lineage contribution^2,3,21^. However, a unifying model that explains the origins and interrelationships of the multiple layers of HSC heterogeneity remains elusive. Here, we performed an in-depth analysis of HSC clonal reconstitution through serial single-cell transplantations, combined with single-cell molecular profiling of clonal systems and mathematical modeling, to establish a unifying framework that clarifies the interrelationships among these distinct layers of molecular and functional heterogeneity.

## Results

### Comprehensive characterization of clonally-derived hematopoietic systems links HSC heterogeneity to reconstitution kinetics

To generate high-resolution maps of HSC clonal heterogeneity, we transplanted single long-term HSCs (phenotypically defined as Lineage-, Kit+, Sca1+, EPCRhi, CD34-, CD150+, CD48-) from GFP expressing mice^22^, along with supportive bone marrow (BM), into 54 lethally irradiated congenic mice (**Fig. 1a**). 6 mice received 30 HSCs each, to act as polyclonal controls. We subsequently performed a detailed kinetics-based analysis of clonal progeny output in recipient mice by interrogating the composition of peripheral blood every four weeks, as well as bone marrow, spleen, lymph nodes, liver, lung, thymus, colon and peritoneal cavity at the 20-week post-transplantation endpoint, utilizing a total of 37 immunophenotypic markers to characterize 55 distinct HSC-derived cell populations. Overall, 34 mice (62.96%) showed donor chimerism >0.1% in any peripheral blood cell type in at least one time point, with 22 (40.74%) demonstrating chimerism above this level at 20 weeks post-transplant (**Supplementary Fig. 1**). Following bulk secondary transplantation of bone marrow from recipients with sustained engraftment, all re-transplanted mice exhibited transient donor chimerism, and 13 out of 18 mice (72.22%) displayed detectable donor-derived cells at the 20-week endpoint. To gain a comprehensive overview of the spectrum of outcomes in recipient mice, we performed principal component analysis (PCA) using the clonal contributions of each transplanted HSC to all measured HSC-derived cell types across all assessed organs at the endpoint (**Fig. 1b-f, Supplementary Fig. 2a-h**). The first dimension distinguished clonal systems with enriched engraftment in hematopoietic stem and progenitor cells (HSPCs), erythroid, megakaryocyte, and myeloid lineages from those which predominantly produced lymphoid cell types (**Fig. 1b,c, Supplementary Fig. 2b,c**). The second dimension broadly segregated clonal systems with high contributions to B versus T cells, while the third dimension separated clonal systems based on their specific chimerism in HSCs, multipotent progenitors, megakaryocyte progenitors and platelets (**Fig. 1e, f, Supplementary Fig. 2e,f**). Notably, separating clonal systems by dimension 1 aligned with a previously proposed classification of long-term repopulating HSCs based on lineage biases^6^. That is, so-called myeloid-biased or “α” HSCs; balanced multilineage or “β” HSCs; and lymphoid-biased or “γδ” HSCs (**Fig. 1d**). Finally, dimension 3 identified previously described platelet-biased HSCs residing at the top of the hematopoietic hierarchy^12^. Based on this analysis, 3 main clusters were identified. Cluster 1 was characterized by high HSPC chimerism in the BM, but relatively low mature hematopoietic cell chimerism, and a strong bias towards platelet and myeloid output; cluster 2 exhibited intermediate HSPC chimerism and a balanced multilineage output; and cluster 3 was characterized by strong lymphoid bias and reduced levels of HSPCs in the BM (**Fig. 1b,e**, **Supplementary Fig. 2h**). In line with previous literature, clonal systems from cluster 1 and 2 showed superior secondary transplantation capacities compared to those of cluster 3 (**Supplementary Fig. 2i**). Importantly, we observed that the heterogeneity between the three clusters appeared to correlate with distinct reconstitution kinetics in the peripheral blood (**Fig. 1g**). Thus, transplanted HSCs within cluster 1 replenished blood cells very slowly with an overall steady increase in chimerism across the window of observation. HSCs within cluster 2 harbored strong engraftment potential and demonstrated more rapid reconstitution kinetics, repopulating up to 75% of all blood cell types after 16 weeks and plateauing around 20 weeks post-transplantation. In contrast, HSCs within cluster 3 engrafted the fastest, but then declined in their blood chimerism from 12 weeks post-transplantation onwards. Collectively, these data confirm previously identified heterogeneity in clonal systems derived from single HSCs with regards to self-renewal capacity and lineage biases, and suggest that these properties might be linked to reconstitution kinetics in a transplant setting^6,8,23^.

**Fig. 1:**
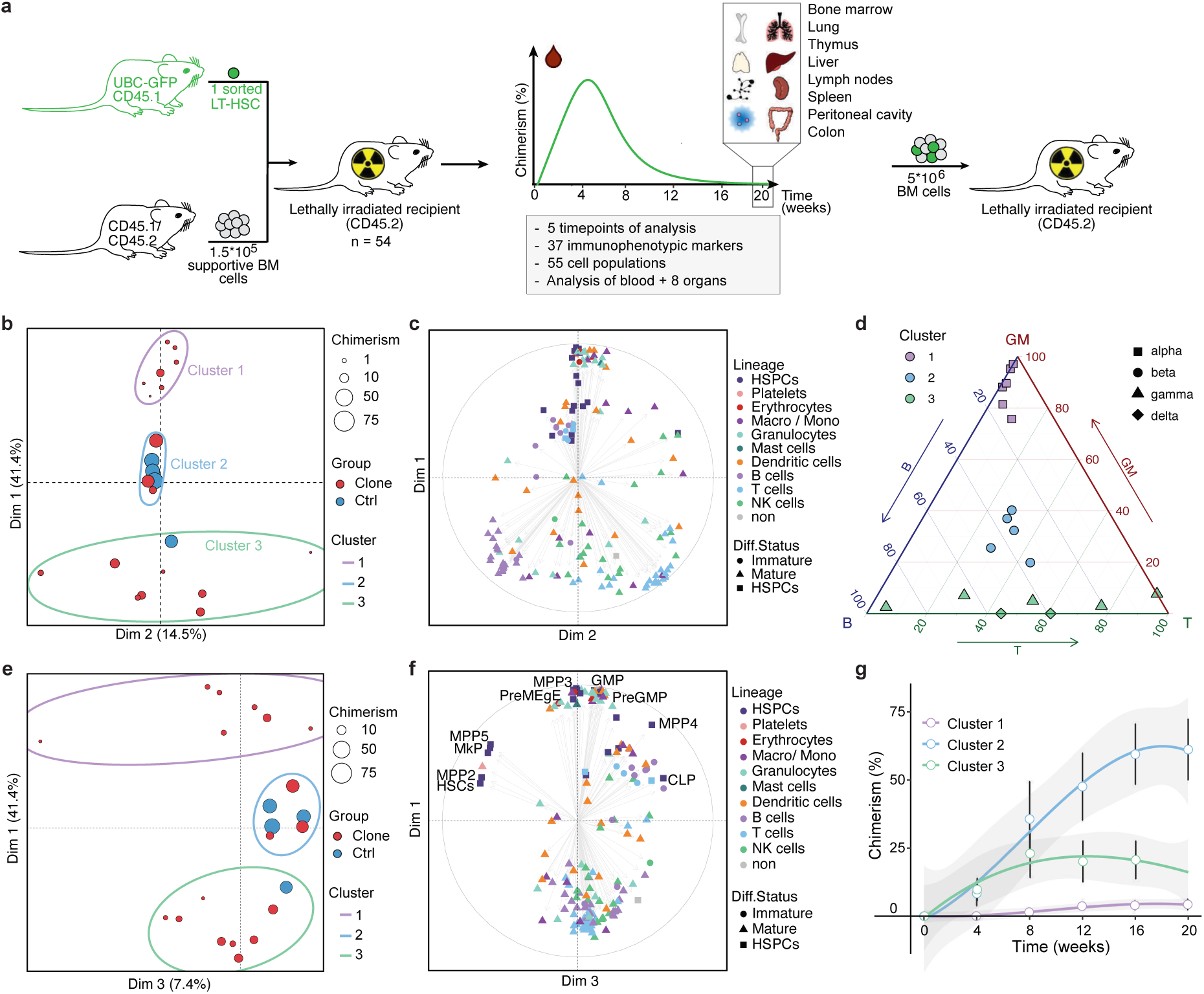
Comprehensive characterization of clonally-derived hematopoietic systems links HSC heterogeneity to reconstitution kinetics. **a)** Schematic overview of experimental design. Single LT-HSCs (or 30 control HSCs) from CD45.1^+^ UBC-GFP donors were transplanted together with 1.5*10^5^ supportive bone marrow cells from CD45.1/2^+^ C57BL/6 mice into lethally irradiated CD45.2^+^ recipient mice. Comprehensive cytometric analysis of donor-derived blood and immune cells was performed every 4 weeks in the blood and across various organs at 20 weeks post-transplant. 5*10^6^ bone marrow cells from primary recipient mice were re-transplanted into secondary recipients to evaluate the repopulation capacity of the clonal systems. **b)** Principal component analysis (PCA) considering the cellular composition of 55 HSC-derived cell types across 9 organs at the endpoint of the primary transplant (week 20). First and second principal components are displayed and overall chimerism is highlighted by dot size. Polyclonal controls (Ctrl) are highlighted in blue. Hematopoietic systems are clustered into three groups based on hierarchical clustering of the top 3 PCs. **c)** Variable contribution map of (b) highlighting the loadings by differentiation status and lineage. **d)** Ternary plot subdividing blood cell repopulation patterns of transplanted HSCs at 20 weeks post-transplant according to GM/(B+T) ratios (encoding symbols), which are calculated dividing the donor chimerism of granulocytes and monocytes by the donor chimerism of B and T cells; as described in^6^. Clonal systems are color-coded based on the clusters annotation from (b) and shaped according to the alpha, beta, gamma, delta classification described in^6^. **e)** First and third principal component projections from PCA described in (b). **f)** Variable contribution map of (e) highlighting the loadings by differentiation status and lineage. **g)** Overall blood cell chimerism over time split between the clusters identified in (b). Dots highlight the mean chimerism and error bars the standard deviation per group and time point. Mean chimerism is smoothly fitted using a third-degree polynomial function with respective confidence intervals highlighted in gray. n = 22 clonal systems analyzed in b-g. Abbreviations: PCA: principal component analysis; UBC-GFP: ubiquitin C-green fluorescent protein; LT-HSC: long-term hematopoietic stem cell; BM: bone marrow; HSPC: hematopoietic stem and progenitor cell; NK cell: natural killer cell; GM: granulocyte/macrophage; Ctrl: control; Dim: dimension; MPP: multipotent progenitor; MkP: megakaryocyte progenitor; PreMegE: pre-megakaryocyte-erythrocyte; GMP: granulocyte-monocyte progenitor; CLP: common lymphoid progenitor.

### A quantitative framework for hematopoietic reconstitution kinetics identifies time-dependent parameters associated with stem cell self-renewal

To further delineate the relationship between reconstitution kinetics and distinct layers of functional HSC heterogeneity, we derived a quantitative framework describing blood reconstitution from HSCs. For this purpose, we modeled the reconstitution kinetics of peripheral blood production by fitting temporal engraftment data for each lineage to a single-humped function (**Fig. 2a**). This function was chosen as it accurately mirrors blood cell reconstitution kinetics, including an initial delay phase, a growth phase, a plateau phase, followed ultimately by a decline phase. This approach allowed us to extract different kinetic-based parameters from the fitted curves, broadly divided into chimerism- and time-dependent parameters, providing quantitative insights into the kinetics of blood cell repopulation (**Fig. 2a, Supplementary Fig. 3a,b**). These chimerism-dependent parameters consist of maximum chimerism level (yMax); overall chimerism (area under the curve, AUC); and the chimerism increase at the initial engraftment phase (slope), while time-dependent parameters include time delay of engraftment (t0); absolute time to reach maximum chimerism (tyMax); time span from initial engraftment until reaching the maximum chimerism (tGrowth); time of decline of chimerism from maximum to half of the initial maximum (tDecline, also referred to as half-life); and the total time taken to transition from initial engraftment, through maximum chimerism, to half of the maximum chimerism (tHalfReg). By introducing a quantitative framework that characterizes each clonal system through these kinetic parameters, broader associations independent of the previously defined clusters could be determined.

**Fig. 2:**
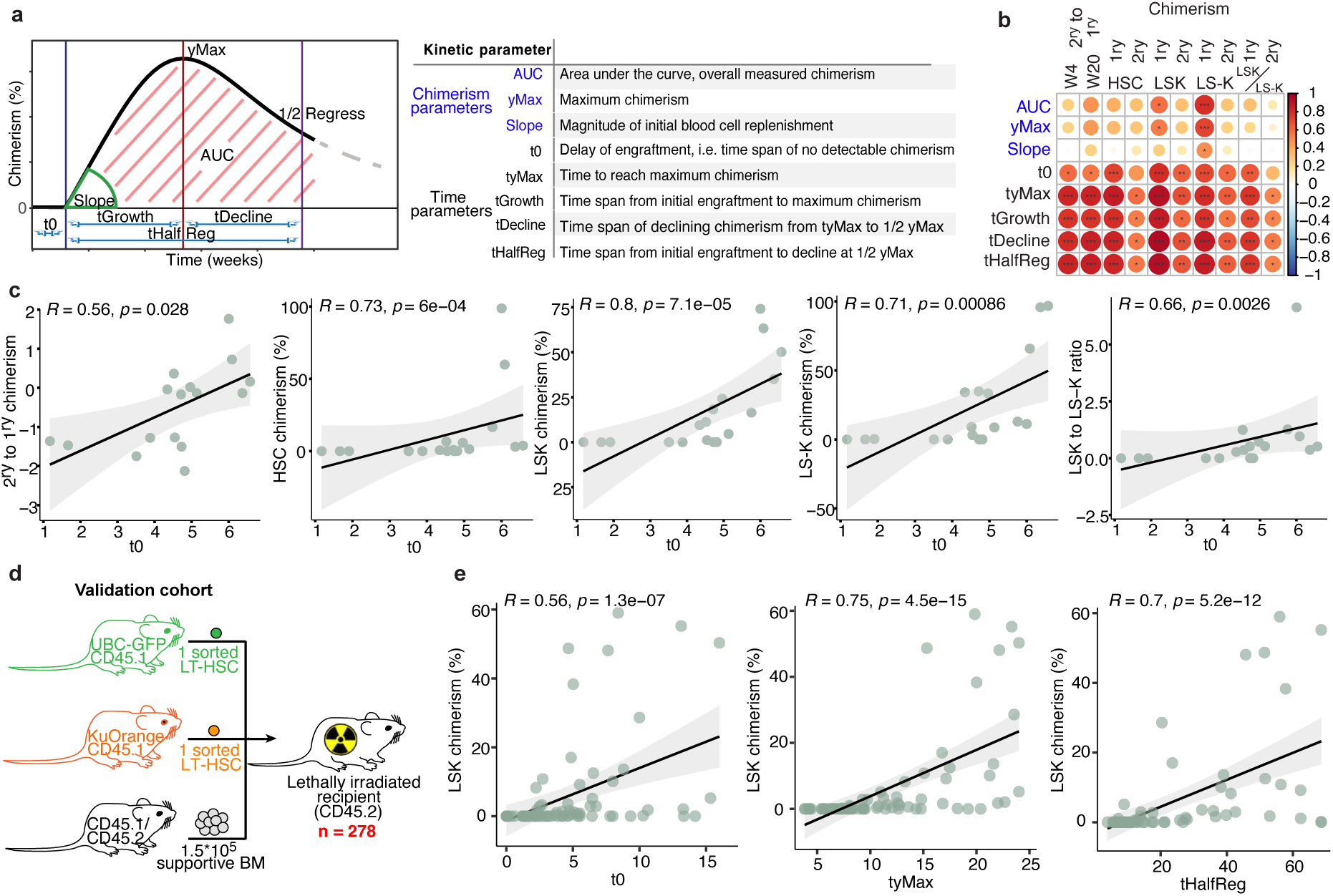
Hematopoietic reconstitution kinetics are linked to functional HSC potency. **a)** Illustration of kinetic parameters. Relative chimerism of each blood cell type per hematopoietic system was fitted using the single humped function and characterized by curve-specific characteristics (kinetic parameters): delay (t0), growth time (tGrowth), decline to ½ max (tDecline), growth and decline time (tHalfReg), maximum chimerism (yMax), time to reach yMax (tyMax), overall chimerism (AUC, area under the curve), steepness of engraftment (slope). **b)** Overview of Spearman correlation analysis between various metrics of HSC functional potency with mean kinetic parameters describing reconstitution kinetics described in (a). Metrics of HSC potency include the difference in peripheral blood chimerism in the secondary bulk transplantation (2ry) compared to the primary (1ry), at 4 (W4) and 20 weeks (W20) post-transplant, HSC chimerism, immature HSPC (LSK) chimerism, committed progenitor (LS-K) chimerism and the ratio of more immature HSPCs (LSK) and committed progenitor (LS-K) at week 20 post-transplant in the primary (1ry) and secondary (2ry) transplantations. Spearman correlation coefficients are displayed by dot size and color. **c)** Exemplary correlation analysis from (b). Correlation between the time delay (t0) of reconstruction and various metrics of HSC functional potency are displayed. Each dot represents a single HSC-derived hematopoietic system. Spearman’s Rho and significance are indicated. **d)** Experimental scheme of co-transplantation of two single LT-HSCs (LSK CD150^+^ CD48^-^ CD34^-^ EPCR^+^) derived from UBC-GFP and KuOrange mouse models, together with 1.5*10^5^ supportive BM cells into lethally irradiated recipient mice. **e)** Spearman correlation analysis between the HSPC chimerism and the kinetic parameters t0, tyMax and tHalfReg. Each dot represents a single HSC-derived hematopoietic system. Spearman’s Rho and significance are highlighted. Significance levels are indicated by: * for p < 0.05, ** for p < 0.01, *** for p < 0.001, **** for p < 0.0001. Abbreviations: PB: peripheral blood; HSC: hematopoietic stem cell; LSK: Lineage-Sca1+cKit+; LS-K: Lineage-Sca1-cKit+; UBC-GFP: ubiquitin C-green fluorescent protein; KuOrange: Kusabira Orange; LT-HSC: long-term hematopoietic stem cell; BM: bone marrow.

To systematically assess associations between HSC functional potency and reconstitution kinetics, we conducted correlation analyses comparing both chimerism- and kinetic-dependent parameters with reconstitution of the peripheral blood, and the primitive hematopoietic stem and progenitor cell (HSPC) compartment in the BM as an indicator of stem cell self-renewal (**Fig. 2b,c**). Conventional chimerism-dependent parameters showed only limited correlation with the degree of regeneration of the HSC and progenitor compartments in the BM. In contrast, all time-dependent kinetic parameters demonstrated a strong correlation with stemness-associated HSPC regeneration (**Fig. 2b,c**). To validate these findings in a much larger independent cohort, we performed an additional 278 single long-term HSC transplants, in which single HSCs isolated from UBC-GFP mice and Kusabira Orange (KuO) mice^24^ were co-transplanted as a pair into recipient mice in order to reduce the total number of required recipient mice (**Fig. 2d**). Overall, 36.44% of the clones showed positive chimerism (>0.1%) at any given time point. Consistent with our initial dataset, we identified kinetics of reconstitution as a variable metric capable of deconvoluting the functional heterogeneity within the transplanted HSC clones (**Supplementary Fig. 4a**). Moreover, dimensionality reduction of the chimerism data was also able to segregate the clonal systems based on their lineage differentiation output and stage of differentiation (**Supplementary Fig. 4b-e**). Using our quantitative framework, we were able to validate the association between time-dependent reconstitution parameters and stemness features (**Fig. 2e**), corroborating the hypothesis that blood reconstitution kinetics are tightly linked to the functional potency of HSCs in a transplant setting.

### Lineage biased HSC output correlates with reconstitution kinetics

Lineage biased output from HSCs has been characterized by the disproportionate production of specific mature cell types and has been linked to stemness characteristics. Specifically, platelet- and myeloid-biased HSCs have been associated with high self-renewal capacities, while lymphoid-biased HSCs are linked to a decline in functional potency^6,7,11^. However, these lineage biases are typically defined by the cellular composition at a single endpoint and do not account for the distinct half-lives of blood cell types or the differing reconstitution kinetics of clonal systems. To explore the relationship between reconstitution kinetics and lineage-biased blood cell production, we first ranked all HSC-derived blood cell types by their first appearance in peripheral blood, as defined by their mean delay parameter t0 (**Supplementary Fig. 3c).** In line with previous reports, platelets were generated first, followed by myeloid cells, and then lymphoid cells^6,7,11^. While this sequence was consistent across most clonal systems, the onset of platelet generation and the delay to the onset of subsequent lineages progressively increased from fast to slow reconstituting clonal systems. Fast clonal systems, with a low t0, showed an early burst in cell type generation across all lineages, followed by a rapid decline in chimerism (**Fig. 3a, Supplementary Fig. 3b,d;** cluster 3 type HSCs). As lymphoid cells exhibit longer half-lives, these systems appear progressively more lymphoid-biased with the passage of time. In contrast, systems with higher t0 appeared myeloid-biased at earlier time points, then progressed to a more balanced mature cell output, sometimes with evidence of an eventual decay of the myeloid lineages at very late time points. Notably, systems with the highest t0 values did not generate myeloid or lymphoid cells during primary transplantation but demonstrated multipotency in secondary transplants. These data link accelerated HSC reconstitution kinetics with a shift from platelet to myeloid and lymphoid output, and suggest that apparent lineage biases may be a function of the time point of analysis post-transplantation rather than representing independent HSC states.

**Fig. 3:**
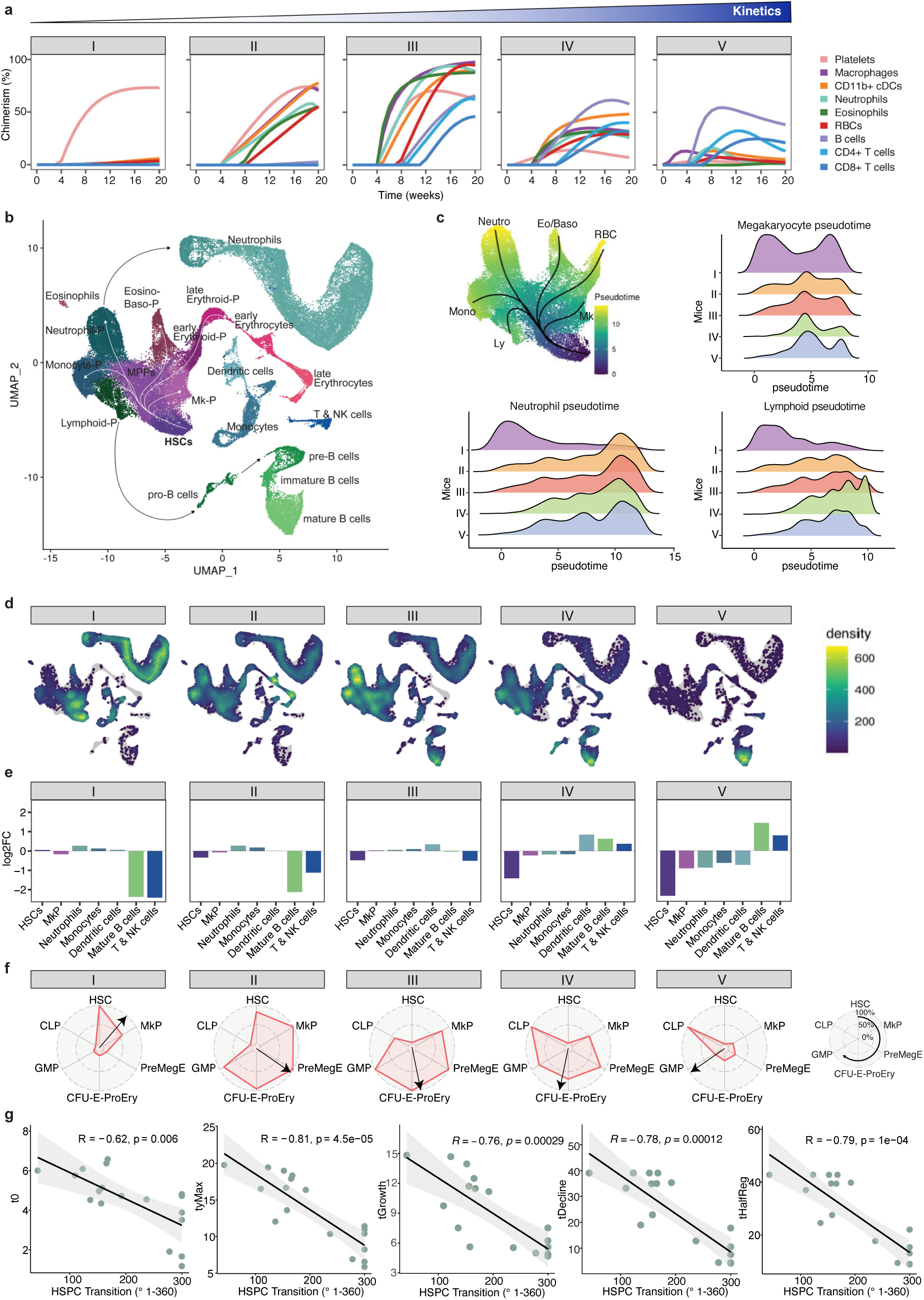
Kinetics of clonal reconstitution is associated with HSC lineage biases. **a)** Reconstitution kinetics of exemplary HSC-derived clonal systems ordered from slow (left) to fast (right). 5 clonal systems selected for single-cell RNA-sequencing (scRNA-seq) are depicted. Respective cell lineages are color-coded. **b)** Global UMAP representation of scRNA-seq data from 20-week bone marrow of the 5 selected clonal, as well as 2 polyclonal, hematopoietic systems from (a). Cell types are highlighted by color. Differentiation trajectories are illustrated by arrows. **c)** Pseudotime inference and density plots visualizing the distribution of cells for each clonally-derived HSPC compartment over pseudotime. Pseudotime from HSC to committed progenitors is color-coded on UMAP and trajectories indicated by arrows (top left). Relative frequencies of differentiation states along pseudotime are indicated for distinct lineages and highlighted separately for each clonal system (I - V) from (a). **d)** Density UMAP representation of the hematopoietic BM ecosystem split by clone from (a) and colored by density and distribution of clonally-derived progeny. **e)** Bar graphs depicting the fold change (log2FC) in clonally-derived hematopoietic cell type abundances from each system, compared to the mean of the polyclonal controls. **f)** Exemplary composition of HSPC compartments of clonally-derived systems I - V, based on the cytometric analysis from Fig. 1. Progenitors are ordered by Pearson correlation distance from HSCs based on clonal compositions of the progenitor compartments 20 weeks post-transplant (see right illustration). The arrow indicates the mean composition of the respective clonally-derived HSPC compartment and illustrates the current state of “HSPC transition”. See Supplementary Fig. 6 for all HSPC compartments. **g)** Spearman correlation analysis of HSPC transition (see (f)) and kinetic parameters of all clonal systems with sustained engraftment from Fig. 1. Each dot represents a single HSC-derived hematopoietic system. Spearman’s Rho and significance are highlighted. Abbreviations: cDC: conventional dendritic cell; RBC: red blood cell; HSC: hematopoietic stem cell; P: progenitor; Mk: megakaryocyte; NK: natural killer cell; Eo: eosinophil; baso: basophil; preMegE: pre-megakaryocyte-erythrocyte; CFU-E-ProEry: colony-forming-unit-erythroid-proerythroblast; GMP: granulocyte-monocyte progenitor; CLP: common lymphoid progenitor; HSPC: hematopoietic stem and progenitor cell.

To gain a deeper understanding of how time-point-resolved lineage-skewed output in the periphery relates to bone marrow hematopoiesis, we performed droplet-based single-cell RNA sequencing (scRNA-seq) on bone marrow progeny from five clonal systems at week 20 post-transplant (**Fig. 3a,b**), representing a spectrum spanning slow to fast reconstitution, as well as two polyclonal controls. This resulted in a clonally-resolved map of 76,863 high-quality cells, covering all major hematopoietic cell types, including differentiation tracks from the most immature HSCs to all lineage-committed progenitors and their continued maturation into blood and immune cells (**Fig. 3b, Supplementary Fig. 5a**). Compositional and trajectory analyses of clonally-derived cell states revealed that lineage-skewed blood production and variable reconstitution kinetics are reflected in the HSPC compartment at the time of harvest (**Fig. 3c-e**). For instance, the slowest reconstituting clone (clone I), which predominantly produced platelets and only began generating myeloid cells by week 20, retained a significant number of progeny in the most primitive HSC and megakaryocyte progenitor (MkP) compartments, with modest occupancy of myeloid progenitor and mature cells. Lymphoid progenitors and maturing lymphoid cells were highly under-represented, reflecting the clonal system’s blood production at that moment in time (**Fig. 3d,e**). In line with this, daughter HSCs derived from slow reconstituting clones displayed transcriptomic signatures associated with low lineage output, high serial engraftment and megakaryocyte bias^8^, while daughter HSCs in faster clones showed a progressive increase in transcriptomic signatures of active multilineage HSCs (**Supplementary Fig. 5b**). Clonal systems with faster reconstitution kinetics demonstrated a progressive shift in abundance of transcriptomically-defined cells. Thus, more rapid reconstitution kinetics correlated with a progressive decrease in the HSC and MkP compartments, accompanied by a transition from systems where the erythro-myeloid lineages dominated at the progenitor and mature cell level, to those where the lymphoid lineages were in the majority (**Fig. 3d,e**). These findings support a model where slowly differentiating HSC clones better regenerate the HSPC compartment, initially produce platelet- and myeloid-skewed progeny, and progressively transition to balanced and lymphoid-biased outcomes as their reconstitution kinetics increase and self-renewal capacity declines.

To validate the link between reconstitution kinetics and lineage biases, we interrogated the clonally-derived HSPC compartments from the deeply immunophenotyped single-cell cohort introduced above (**Fig. 1**). By ordering lineage-committed progenitors clockwise based on their Pearson correlation distance to HSCs, we generated clock-like representations of the HSPC compartment (**Fig. 3f, right**). Upon arranging all 22 clonal systems with sustained engraftment along this "clock" framework, based on their mean HSPC composition - termed “HSPC transition time” - we were able to characterize systems ranging from those predominantly retaining HSCs (positioned closer to 12 o’clock) to those with progressively more differentiated HSPC phenotypes (**Fig. 3f, Supplementary Fig. 6**). Consistent with our previous findings, this analysis revealed a strong association between HSPC transition time and lineage biased output (**Fig. 3g**). That is, slowly reconstituting systems retained a highly immature and megakaryocyte-primed HSPC compartment correlating with a restriction to platelet and myeloid generation, while faster systems showed a progressive shift to myeloid and lymphoid-primed HSPCs as production of mature cells skewed to balanced and then lymphoid outcomes. Through this transition, the primitive HSC compartment is progressively exhausted. Overall, our findings suggest that conventional categorical definitions of lineage biases and stemness are highly dependent on the time point of investigation and the underlying kinetics of the clonal system. In contrast, kinetics-based parameters provide an alternative approach for classifying clonal hematopoietic systems in a continuous manner.

### Single-cell re-transplantation of clonally-derived daughter HSCs demonstrate a unidirectional transition from slow to fast reconstitution kinetics

Our data infer a hierarchical relationship between slow and fast-engrafting HSC clones, where slow engrafting clones would be more primitive and potentially the precursor of clones with faster reconstitution kinetics. To investigate this hypothesis, we developed a mathematical model of the process and tested it using our time-resolved chimerism data. Initially, we categorized clonal systems as either fast- or slow-engrafting based on their blood reconstitution kinetics. We then assessed whether this dichotomy could be explained by a linear hierarchy within the HSC compartment, consisting of an upstream (slow) and downstream (fast) sub-compartment (**Fig. 4a**). The model successfully reproduced the sequential production order of blood cell types post-transplantation and captured the lineage biases associated with the distinct kinetics of blood production (**Fig. 4b**). Specifically, the slow system resulted from transplanted HSCs populating the upstream compartment, while the fast system was driven by HSCs populating predominantly the downstream compartment. These findings suggest that the experimental data fit a model describing transitions from slow- to fast-reconstituting clones in a linear hierarchy, associated with distinct kinetics of lineage contributions and declining functional potency. To experimentally validate this model, we performed serial single-cell transplantations, so that the post-engraftment output of individual daughter HSCs could be directly compared to that of the parent HSC. Such comparisons cannot be drawn by HSC barcoding approaches, since all daughter HSCs of a barcoded HSC will share the exact same barcode and will therefore be indistinguishable from each other. We selected six primary recipients of single HSCs which showed robust chimerism in the HSPC compartment at the 24-week post-transplantation experimental endpoint and which demonstrated slow- to intermediate-reconstitution kinetics. We harvested single HSCs from these donors and re-transplanted a total of 525 single daughter cells into secondary recipients, representing the majority of the HSC reserve that we could purify from the primary recipients (**Fig. 4c**). The overall percentage of daughter HSCs with detectable engraftment in peripheral blood at any time point declined from 36.3% in primary recipients to 9.9%, in secondary recipients, while the percentage of clones with long-term engraftment capacity dropped even more dramatically, from 24.8% to 2.6%, (**Fig. 4d,e**). In line with this observation, the reconstitution of the HSPC compartment by daughter HSCs declined significantly in the majority of clones (**Fig. 4f**). Remarkably, all but one of the re-transplanted HSCs (99.8%) exhibited decreased chimerism levels compared to their parent HSC at week 24 post-transplant (**Fig. 4e**), representing a decline in functional potency in virtually all secondary HSC clones and suggesting that full self-renewal is a very rare event in the context of transplantation. To investigate whether these data are consistent with the kinetic hierarchy model (**Fig. 4a, b**), we quantified kinetic parameters for both parent- and daughter-derived clonal systems. Compared to their respective parents, daughter clonal systems exhibited significantly accelerated kinetics across all parameters (**Fig. 4h, Supplementary Fig. 7**). Notably, almost all daughter stem cells unidirectionally shifted from myeloid-biased to more lymphoid-biased blood production, consistent with our previous data linking faster reconstitution kinetics to this shift in lineage output (**Fig. 4g**). In very rare cases, daughter clones maintained reconstitution kinetics comparable to their parents, which was linked to high self-renewal of the HSPC compartment and the maintenance of myeloid-to-lymphoid ratio in secondary transplantation endpoint analyses (**Fig. 4f-h, Supplementary Fig. 7**). These findings support the prediction from our model that slow engrafting clones unidirectionally give rise to faster engrafting HSC clones, associated with a progressive change in lineage output and loss of functional potency.

**Fig. 4:**
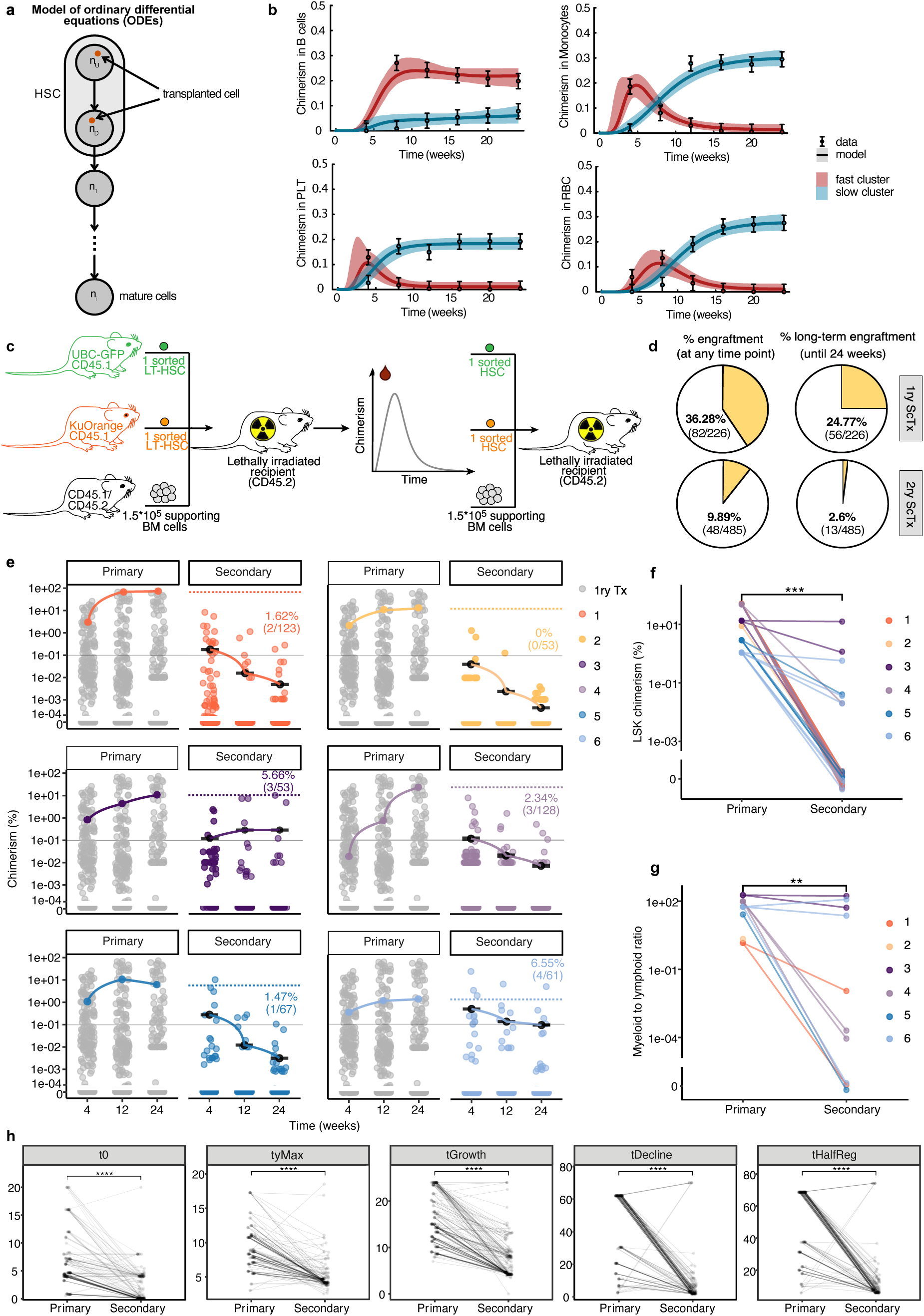
A shift from slow to fast kinetics in daughter HSCs is associated with a decline in functional potency. **a)** Illustration of mathematical model in which HSCs can initiate differentiation from an upstream or downstream compartment. **b)** Average chimerism data in peripheral blood cells from fast and slow reconstituting clones (experimental data, validation cohort) and fits derived from the mathematical model described in (a) are displayed. n = 8 clones per cluster **c)** Experimental scheme of secondary single-cell transplantations. **d)** Pie charts displaying the percentage of clones with successful engraftment (>0.1% chimerism in peripheral blood) at any time point post-transplantation (left), and with positive long-term chimerism (at 24 weeks post-transplantation) (right), for primary (top) and secondary (bottom) transplants. 1ry ScTx: primary single-cell transplantation; 2ry ScTx: secondary single-cell transplantation. **e)** Chimerism in peripheral blood at 4, 12 and 24 weeks post-transplantation comparing each primary to their corresponding secondary daughter HSC transplant. Within each primary plot, the chimerism of the respective parent HSC is highlighted by color. Dotted lines in the secondary plots indicate the maximum blood chimerism reached by the parent HSC in the primary transplantation. The percentage and fraction of long-term engrafting clones is indicated at the 24-week time point of each secondary transplantation plot. n = 6 paired primary to secondary transplantations, each of them with n = 60-141 single transplanted HSCs. **f)** Percentage of Lineage-, Sca1+, Kit+ HSPC (LSK) chimerism in the bone marrow at 24 weeks post-transplantation comparing the primary and secondary transplantations. Each number corresponds to a paired analysis between an individual parent HSC and its corresponding daughter HSCs. n = 51 clonal systems. **g)** Ratio of myeloid to lymphoid progeny of primary and secondary daughter HSC transplantation measured at 24 weeks post-transplantation in peripheral blood. Each number corresponds to a paired analysis between an individual parent HSC and its corresponding daughter HSCs. n = 13 clonal systems **h)** Reconstitution parameters (t0, tyMax, tGrowth, tDecline, tHalfReg) in primary and corresponding secondary single-cell transplantations. Significance was tested by paired Wilcoxon test and is indicated as follows: * for p < 0.05, ** for p < 0.01, *** for p < 0.001, **** for p < 0.0001. Abbreviations: HSC: hematopoietic stem cell; PLT: platelet; RBC: red blood cell; UBC-GFP: ubiquitin C-green fluorescent protein; KuOrange: Kusabira Orange; LT-HSC: long-term hematopoietic stem cell; BM: bone marrow; LSK: Lineage-Sca1+cKit+.

### Cellular competition between progeny of slow and fast engrafting clones contributes to extrinsic regulation of lineage-biased HSC output

Our paired daughter cell experiments support the concept of cell intrinsic inheritance of functional properties, as we observe a generational acceleration in engraftment kinetics alongside decreased self-renewal. However, in our studies, single purified HSCs are not transplanted in isolation; rather, they are co-transplanted with additional supportive bone marrow. This indicates that total blood cell production arises from the combined output of both slow- and fast-reconstituting clones, suggesting that these co-existing clonal systems may interact to regulate their overall output. To explore this concept, we exploited the fact that both the total number of long-term engrafting HSCs (at 24 weeks) and their mature progeny vary from mouse to mouse. We employed mathematical modeling to simulate this process, devising two mutually exclusive models: Model 1 lacks feedback regulation, while Model 2 incorporates feedback regulation where the sizes for mature cell compartments are set by negative feedback (**Fig. 5a**). As expected, in Model 1 the variation in total HSC numbers, as measured by the coefficient of variation, were passed on to the variation in mature cell numbers, whereas in Model 2 feedback regulation strongly reduced the variation in the cellularity of mature blood populations (**Fig. 5b)**. This suggests that, despite variability in contributions from individual HSC-derived hematopoietic systems, the production of mature hematopoietic cells remains tightly constrained by homeostatic feedback mechanisms that enforce strict compartment size limits (**Supplementary Fig. 8a**). In contrast, HSCs exhibited a significantly higher coefficient of variation, suggesting that the compartment size limit, if present at all, is much weaker for HSCs or has not been reached in the setting where only a small number of input stem cells have been transplanted. Given the compartment size restriction in mature cell populations, we reasoned that the replenishment of mature blood populations by single HSC clones might be influenced by competing HSC clones. If correct, this hypothesis would predict that fast engrafting clones would rapidly fill up cellular compartments to their limit, while slowly engrafting clones would only be able to contribute to mature cell production once the levels of mature blood cells had declined below this limit, due to exhaustion of fast engrafting HSCs and their progeny. To investigate this hypothesis, we quantified the absolute levels of mature blood cells produced by the transplanted single HSCs versus those derived from the co-transplanted supportive bone marrow in the same mice. Consistent with our hypothesis, we observed a strict inverse correlation between absolute numbers of mature cells generated by the single LT-HSCs and the co-transplanted supporting bone marrow (**Supplementary Fig. 8b**). Notably, the competition between clonal offspring and supportive bone marrow was low at the beginning of transplantation and gradually increased over time, eventually plateauing as the compartments reached their maximum compartment size limit (**Supplementary Fig. 8c**). Collectively, these data suggest that feedback regulation restricts the compartment size of mature blood populations, and raise the possibility that clonal competition between slow- and fast-reconstituting clones might contribute to lineage biases in a cell-extrinsic manner. Thus, we hypothesized that after the exhaustion of the HSPC compartment in fast engrafting clones, the myelo-erythroid output will decline more rapidly than the much longer-lived lymphoid lineages. Therefore, more slowly engrafting clones will initially only have space to produce myelo-erythroid progeny, while new lymphoid progeny could only be generated later, once the lymphoid progeny of fast engrafting clones had declined to the extent that they failed to sustain the compartment size limit for these lineages (see model: **Supplementary Fig. 8d**).

**Fig. 5:**
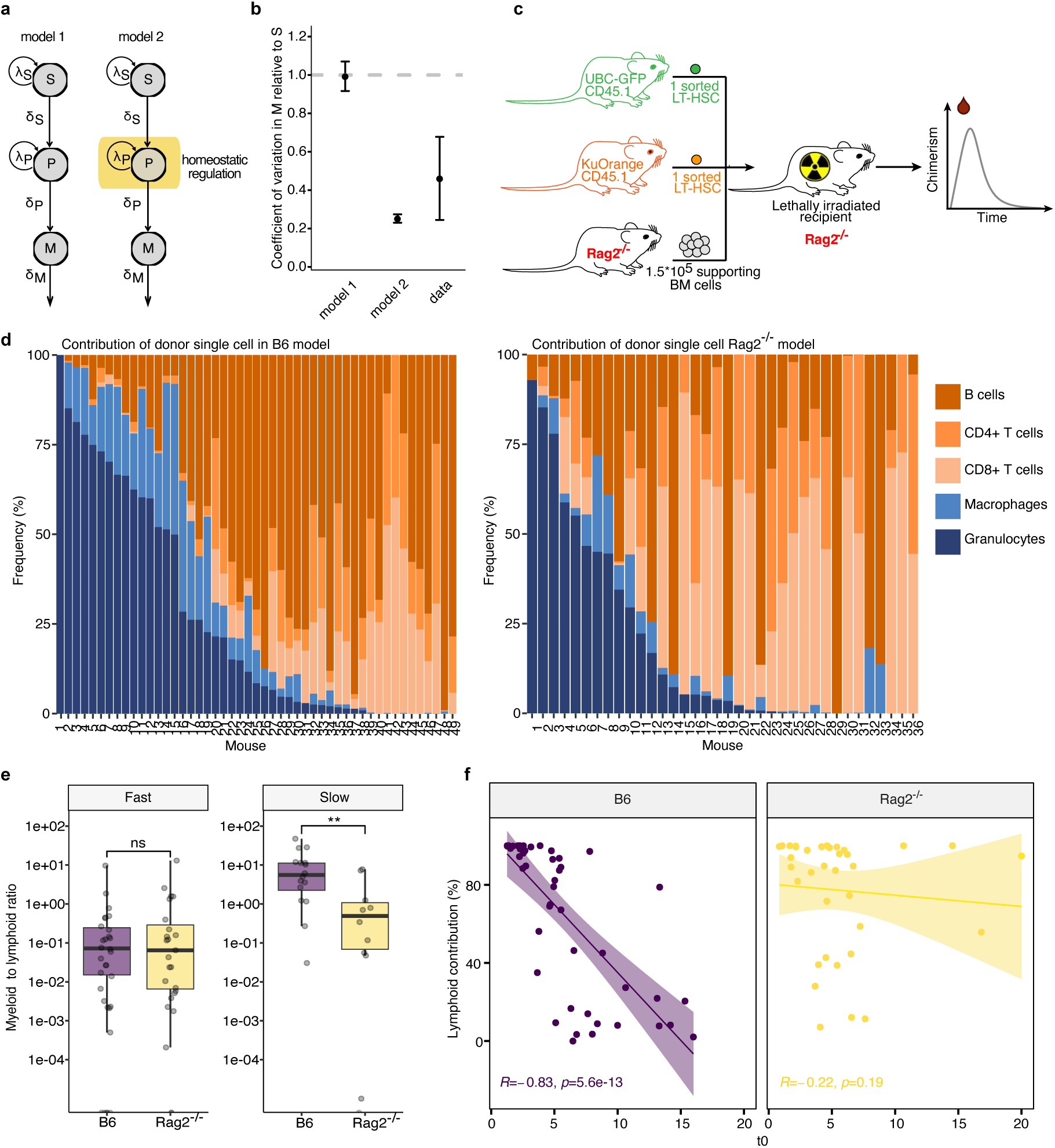
Cellular competition contributes to the establishment of HSC lineage biases. **a)** Illustration of mathematical models describing the process of HSC hematopoietic differentiation in the absence (model 1) or presence of homeostatic regulation (model 2). **b)** Coefficient of variation in mature cells (M) relative to stem cells (S) predicted by model 1 and 2 from (a), compared to experimental data. n = 129 clones. **c)** Experimental scheme of single-cell transplantations in Rag2^-/-^ recipients using Rag2^-/-^ supportive bone marrow cells. **d)** Frequency of indicated cell types produced by the single HSC at 24 weeks post-transplantation in peripheral blood using regular C57BL/6 (B6) supportive bone marrow and recipients (left), or using Rag2^-/-^ supportive bone marrow and recipients (right). **e)** Ratio of myeloid to lymphoid frequency in peripheral blood at 24 weeks post-transplantation, comparing fast (t0 < 6)- and slow (t0 > 6)-reconstituting clones transplanted in the B6 or Rag2^-/-^ systems. **f)** Spearman correlation analysis between the average time delay (t0) in reconstitution of a single HSC and its percentage of lymphoid contribution in peripheral blood at 24 weeks post-transplantation, in B6 versus Rag2^-/-^ hosts. Spearman’s Rho and significance are indicated. n = 36-49 Rag2^-/-^ or B6 clonal systems used in d-f. If not stated otherwise, significant differences between groups were tested by a two-sided Wilcoxon rank-sum test. Significance levels are indicated by: ns for not significant, * for p < 0.05, ** for p < 0.01, *** for p < 0.001, **** for p < 0.0001. Abbreviations: S: stem cell; P: progenitor; M: mature cell; λ: proliferation rate; δ: differentiation rate; UBC-GFP: ubiquitin C-green fluorescent protein; KuOrange: Kusabira Orange; Rag2^-/-^: homozygous knock-out of the recombination activating gene 2; BM: bone marrow; B6: C57BL/6J mouse model.

To interrogate this hypothesis, we made use of the congenic *Rag2* knockout mouse (*Rag2*^-/-^) model^25^, whose HSCs are capable of erythro-myeloid reconstitution but lack the ability to produce mature lymphoid cells (**Fig. 5c**). We transplanted a total of 154 single *Rag2*-wild type UBC-GFP or KuO LT-HSCs, together with *Rag2*^-/-^ supporting bone marrow cells, into *Rag2*^-/-^ recipient mice. In this setting, neither the supporting bone marrow nor the residual recipient hematopoiesis could contribute towards filling the mature lymphoid compartment size limit. We then measured the clonal lineage output and reconstitution kinetics of the wild type HSCs and compared this to wild type LT-HSCs transplanted into wild type recipients along with wild type supporting bone marrow (**Fig. 5c,d**). Interestingly, we observed both fast and slow engrafting clones in both experimental arms, suggesting that kinetic parameters are independent of the lineage output of the co-transplanted competitor cells. However, slow engrafting clones demonstrated an altered lineage output dependent on the cell extrinsic environment (**Fig. 5e**). While slow clones co-transplanted with lymphoid-proficient competitors displayed a pronounced myeloid bias, those transplanted into a *Rag2*^-/-^ hematopoietic system did not exhibit the same lineage skewing. These findings demonstrate that the link between kinetic-based reconstitution parameters and apparent myeloid bias can be uncoupled by modulating the capacity of competitor HSPCs to contribute towards filling mature lymphoid lineages to their compartment size limits. (**Fig. 5f, Supplementary, Fig. 8e**). Taken together, these data provide compelling evidence that apparent intrinsic lineage biases are in fact highly dependent on cell extrinsic regulation, resulting from a competition between slow and fast engrafting HSC clones to saturate the production of mature blood cells until lineage-specific compartment sizes are filled.

## Discussion

Numerous studies have used single-cell transplantations to describe functional heterogeneity within the primitive HSC pool^5–7,10,26–28^, including at the level of reconstitution kinetics^23^. However, there has been a lack of clarity regarding whether these diverse phenotypic outcomes represent discrete intermediates in a branched hierarchy of the most primitive HSCs, or rather cell states aligned along a linear trajectory. This has been particularly confusing with regards to the relationship between HSC lineage bias and multipotency, since the concept of progressive lineage commitment does not seem to be compatible with a model where the most primitive HSCs demonstrate an intrinsic lineage bias, yet generate HSC progeny which are more permissive in the spectrum of cell types they can produce. Our alignment of functionally heterogeneous outcomes along a kinetics-based scale not only provides a novel framework to assign potency based on self-renewal capacity, but also offers a new rationale to explain why the most potent HSCs predominantly produce platelet and myeloid progeny once they first contribute to mature blood cell production, despite being multipotent^11–13^. Indeed, the revelation that lineage-skewed output from slower engrafting clones is likely driven by extrinsic feedback from the mature progeny of more rapidly engrafting competitor clones has important implications for our understanding of normal and diseased hematopoiesis and perhaps also explains why data demonstrating a concrete molecular basis for such biases in the HSC compartment has not yet emerged.

A kinetics-based functional hierarchy aligns well with other transplantation-based studies that clearly support successive waves of HSC clones contributing to mature blood cell production where sustained engraftment and regeneration of the HSC pool was supported by slow or low output clones, including barcoding approaches in murine and primate HSCs^8,29^ and analyses of human engraftment based on retroviral integration sites^30,31^. However, it remains unclear how this relates to the setting of native hematopoiesis, where the transition time from primitive HSCs through to mature blood cells is longer and challenging to measure in an experimental setting in the absence of stimuli that provoke emergency hematopoiesis^32,33^. Nonetheless, it is tempting to speculate that slow-engrafting HSC clones may equate to so-called dormant HSCs, which maintain a state of long-term quiescence during native unperturbed hematopoiesis^28^. Certainly, both cell types appear to represent a subset of highly potent HSCs which have an inherent capacity to restrict their output of progeny, either in the face of pro-proliferative stimuli acting over the course of long time periods in the native niche, or in a myeloablated niche. A direct comparative analysis is restricted by the fact that both cell types can only be identified retrospectively, but it would be interesting to understand the underlying molecular basis for this restricted output, as well as how and why such HSCs eventually overcome this restriction following a temporal delay.

One setting of native hematopoiesis where our findings may be of immediate relevance is the accumulation of myeloid-biased HSCs during aging, which has been attributed as the root cause of a number of age-associated pathological processes ranging from the evolution of myeloid malignancies to immune dysfunction^34–36^. One could extrapolate from our data that aging may result in a progressive accumulation of multipotent HSCs with delayed kinetics and therefore appear myeloid-biased following transplantation. Perhaps such a phenotype might even be selected for during aging, since clones that actively contribute to blood formation will be preferentially lost from the HSC pool^37,38^. This hypothesis aligns with the enrichment of so-called latent HSCs within aged murine bone marrow, which demonstrate low output myeloid-skewed production in primary recipients, but give rise to robust multilineage reconstitution upon secondary transplantation^27^.

Collectively, our study identifies reconstitution kinetics as a unifying metric for classifying primitive HSCs according to their functional potential and provides a novel underlying rationale for lineage-skewed output from these multipotent cells. Furthermore, the kinetics-based principles outlined in this manuscript may have broad relevance for understanding the establishment and remodeling of clonal mosaicism during the development and aging of other regenerating tissues throughout the body.

## Acknowledgements

We would like to thank the DKFZ Flow Cytometry Core facility, the DKFZ center for preclinical research, the DKFZ Next Generation Sequencing Core facility, the Omics IT and Data Management Core facility, and the Single Cell Open Lab. E.R.-C. was funded by the German-Israeli Helmholtz International Research School “Cancer-TRAX”. R.M. and V.K. were supported by the Helmholtz Program Natural, Artificial and Cognitive Information Processing (NACIP) and the Helmholtz Information and Data Science School for Health (HIDSS4Health). N.A. and S.L. were supported by the DKFZ International Postdoctoral Fellowship Program. S.H. received support by the Heisenberg program of the German Research Foundation (DFG), the e:Med LeukoSyStem consortium (BMBF), the HEROES-AYA consortium (BMBF), the TEP-CC consortium (Bruno and Helene Jöster Foundation), the CRC1588 project C03, and the DFG project HA 8790/3-1. This project is co-funded by the European Union (ERC, InteractOmics, 101078713 to S.H.). Views and opinions expressed are, however, those of the author(s) only and do not necessarily reflect those of the European Union or the European Research Council. Neither the European Union nor the granting authority can be held responsible for them. We also wish to thank the Dietmar Hopp Foundation for their generous support. Schematic overview was generated using BioRender.com. We would like to thank Alejo Rodríguez Fraticelli for advice and critical appraisal of this work.

## Author contributions

E.R.-C., F.G., N.A., J.A.-S, M.D.M., S.H., and D.H designed and directed the experimental scheme of work. E.R.-C., F.G., N.A., J.A-S, M.B., F.F., J.J., I.G., J.K., M.D., T.A., M.B.-S., S.L., P.H.-M., J.G. and D.V. performed experiments. F.G., E.R.-C., A.S., A.M., S.H., D.H., M.D.M., A.T. and J.Z. carried out data analysis and/or interpretation of experimental data. T.N., C.L. and V.K. performed the mathematical modeling with help from F.G. and supervision from T.H. and R.M.. S.H., M.D.M., D.H., E.R.-C., and F.G. generated the figures and wrote the manuscript.

## Materials and Methods

### Animal experiments

All animal experiments were approved by the Animal Care and Use Committees of the German Regierungspräsidium Karlsruhe für Tierschutz und Arzneimittelüberwachung (Karlsruhe, Germany) under the TVAs G-41/19 and G-50/17. Mice were maintained in individually ventilated cages under specific pathogen-free (SPF) conditions at the German Cancer Research Center (DKFZ, Heidelberg). Wild type mice (C57BL/6J) were obtained from Janvier Laboratories. Recipient mice were 8-12 weeks when experiments were initiated. UBC-GFP and KuOrange (KuO) mice were used as donors for transplantation experiments. The *Rag2*^-/-^ mouse line was used for recipient mice and to isolate supportive bone marrow.

### Single-cell transplantations

CD45.2^+^ C57BL/6J mice were lethally irradiated with two rounds of 500 Rad. 24 hours later, mice were transplanted via *i.v.* injection with a single CD45.1^+^ LT-HSC (EPCR^hi^, CD34^-^, CD150^+^, CD48^-^, LSK) derived from a transgenic CD45.1^+^ UBC-GFP donor mouse, together with 1.5*10^5^ WT CD45.1^+^/CD45.2^+^ supportive whole bone marrow cells. In a second group of experiments, co-transplantations of single CD45.1^+^ UBC-GFP LT-HSC plus single CD45.1^+^ KuO LT-HSC together with 1.5*10^5^ WT CD45.1^+^/CD45.2^+^ supportive whole bone marrow cells were performed. Co-transplantation were also performed in combination with *Rag2*^-/-^ supportive bone marrow into *Rag2*^-/-^ recipient mice. Engraftment potential was assessed at 4, 8, 12, 16, 20 and in some cases at 24 weeks post-transplantation in peripheral blood cells, and at 20 or 24 weeks in the bone marrow. The discovery cohort also included chimerism analysis in spleen, lymph nodes, liver, lung, thymus, colon and peritoneal cavity at 20 weeks post-transplant. Secondary engraftment potential was evaluated by re-transplanting 5*10^6^ total bone marrow cells or by single-cell transplantations of donor-derived HSCs (GFP^+^ or KuO^+^) from primary recipient mice.

### Bleeding and hematopoietic cell isolation

Peripheral blood was withdrawn from the *vena facialis* and collected into EDTA-coated tubes. Blood cell counts were analyzed using a Hemavet 950 FS (Drew Scientific) or ScilVet abc-Plus+ veterinary blood cell counting machine (Scil GmbH). For the comprehensive immunophenotypic characterization, hematopoietic cells were collected from the peritoneal cavity (PerCav) in 2 mL PBS, and hematopoietic organs and tissues were dissected, including bones, spleen, lymph nodes (LNs), thymus, lung and liver. Bone marrow (BM) was harvested by isolating, cleaning and crushing the vertebral column, tibia, femur, limbs and sternum of sacrificed mice in RPMI + 2% FCS. Cell suspensions were filtered through a 40 μm cell strainer, centrifuged and resuspended in ACK buffer for red blood cell lysis for 3 minutes at room temperature (RT). After washing, 5*10^6^ BM cells were used for secondary transplantation, 3*10^7^ cells were kept for subsequent flow cytometric analysis and the remaining BM was used for scRNA-sequencing and stored in liquid nitrogen until further use. Lungs and liver were minced into small pieces. Lungs were further filtered initially through a 100 μm and subsequently through a 70 μm cell strainer. Liver, LNs, spleen and thymus were filtered through a 40 μm cell strainer. Cell suspensions were spun down, resuspended in RPMI + 2% FCS and split for multiple flow cytometric analysis. Colons were turned inside out, cleaned and incubated in 25 mL extraction medium (RPMI 1640 + 2% FCS + 1 mM DTT + 0,5 mM EDTA) for 20 min at 37 °C to digest the intraepithelial layer. 1 mL FCS was then added to block the digestion, samples were filtered through a 40 μm cell strainer, centrifuged and resuspended in RPMI + 2% FCS for staining. If not stated otherwise, each step was performed on ice, RPMI or PBS supplemented with 2% FCS was used for washing and resuspending and centrifugation was done at 600 g, 4°C for 5 min. For the large-scale validation cohort, the same experimental protocol was followed for the isolation of bone marrow hematopoietic cells.

### Isolation of murine EPCR^hi^ LT-HSCs cells via FACS

Bone marrow cell suspension was subjected to depletion of mature blood cell lineages incubating with a mix of rat anti-mouse biotin-conjugated lineage markers (4.2 µg/mL CD5, 4.2 µg/mL CD8a, 2.4 µg/mL CD11b, 2.8 µg/mL B220, 2.4 µg/mL Gr-1, 2.6 µg/mL Ter-119) for 40 min at 4°C. After incubation, cells were washed once with PBS + 2% FBS, span down at 350 g at 4°C for 5 min, resuspended in 800 µL PBS + 2% FBS and mixed with 800 µL Biotin Binder Dynabeads (Thermo Fisher), which were previously washed (2 washes with PBS + 2% FBS). Beads were added at a concentration of 1 mL beads /1*10^8^ cells. Cells-beads mix was incubated for 45 min at 4°C with constant rotation. Subsequently, lineage-positive cells were depleted using a magnetic particle concentrator (Dynal MPC-6, Invitrogen), and the resulting LSK-enriched fraction was washed once with PBS + 2% FBS, and stained with the panel of antibodies indicated in **Table 1a,b** for 30 minutes at 4°C. After the incubation, stained cells were washed once with PBS + 2% FBS, resuspended in a final concentration of 2 mL PBS + 2% FBS and filtered through a 40 µm cell strainer FACS tube before the sort. All sorting experiments were performed using a BD FacsAria I or II flow cytometer (BD Bioscience) with a 100 μm nozzle and single-cell purity. Single EPCR^hi^ LT-HSCs (**Supplementary Fig. 9**) were sorted into round-bottom 96-well plates with 100 μL RPMI + 2% FBS with a cooling system. After the sort, 100 µL of supportive total bone marrow at a concentration of 1.5*10^6^ cells/ mL was added in each well on top of the sorted single HSC using a multichannel pipette, reaching a final volume of 200 µL per well.

**Table 1a.**
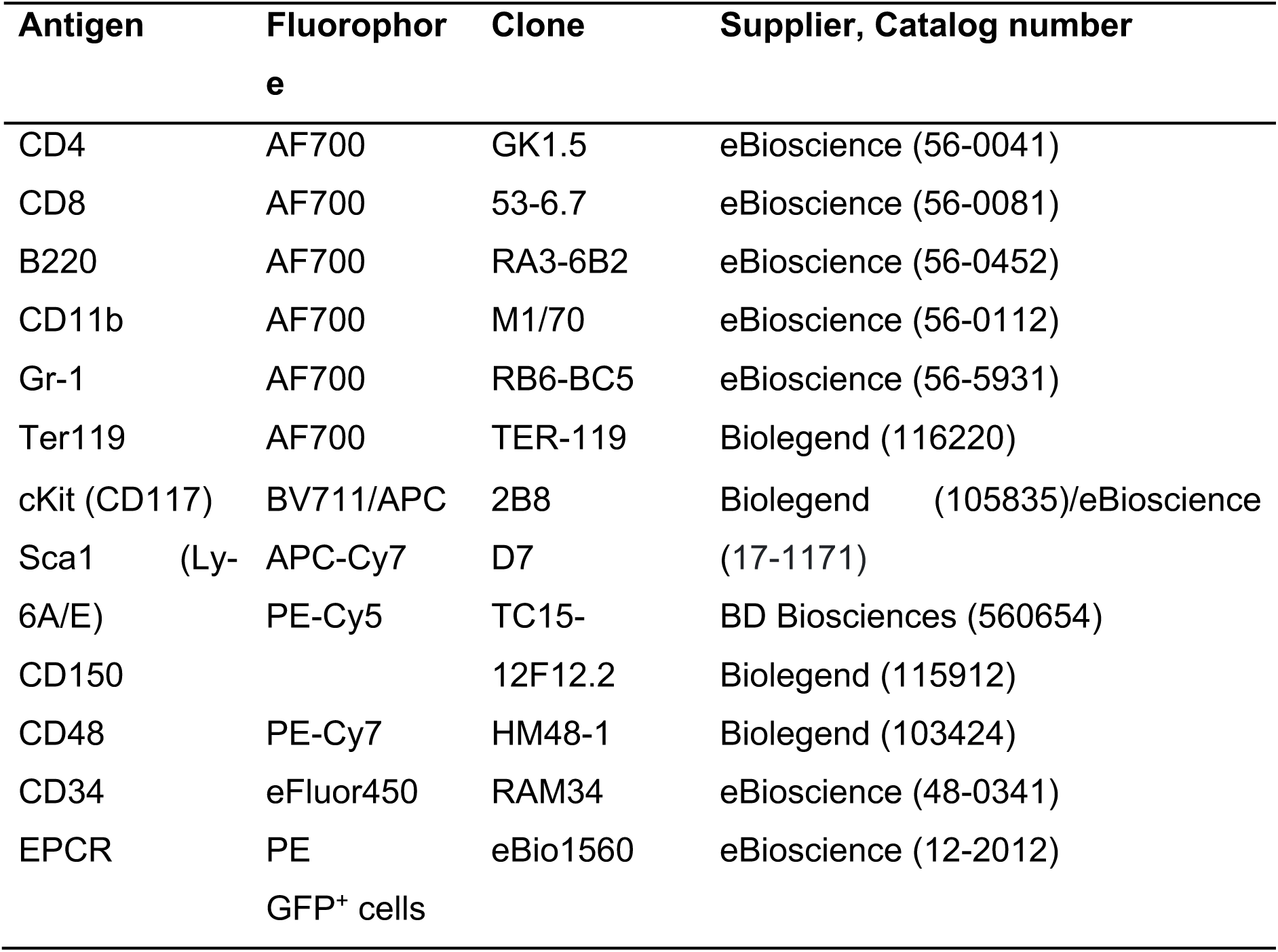
Antibody panel for the isolation of GFP^+^ EPCR^hi^ LT-HSCs.

**Table 1b.**
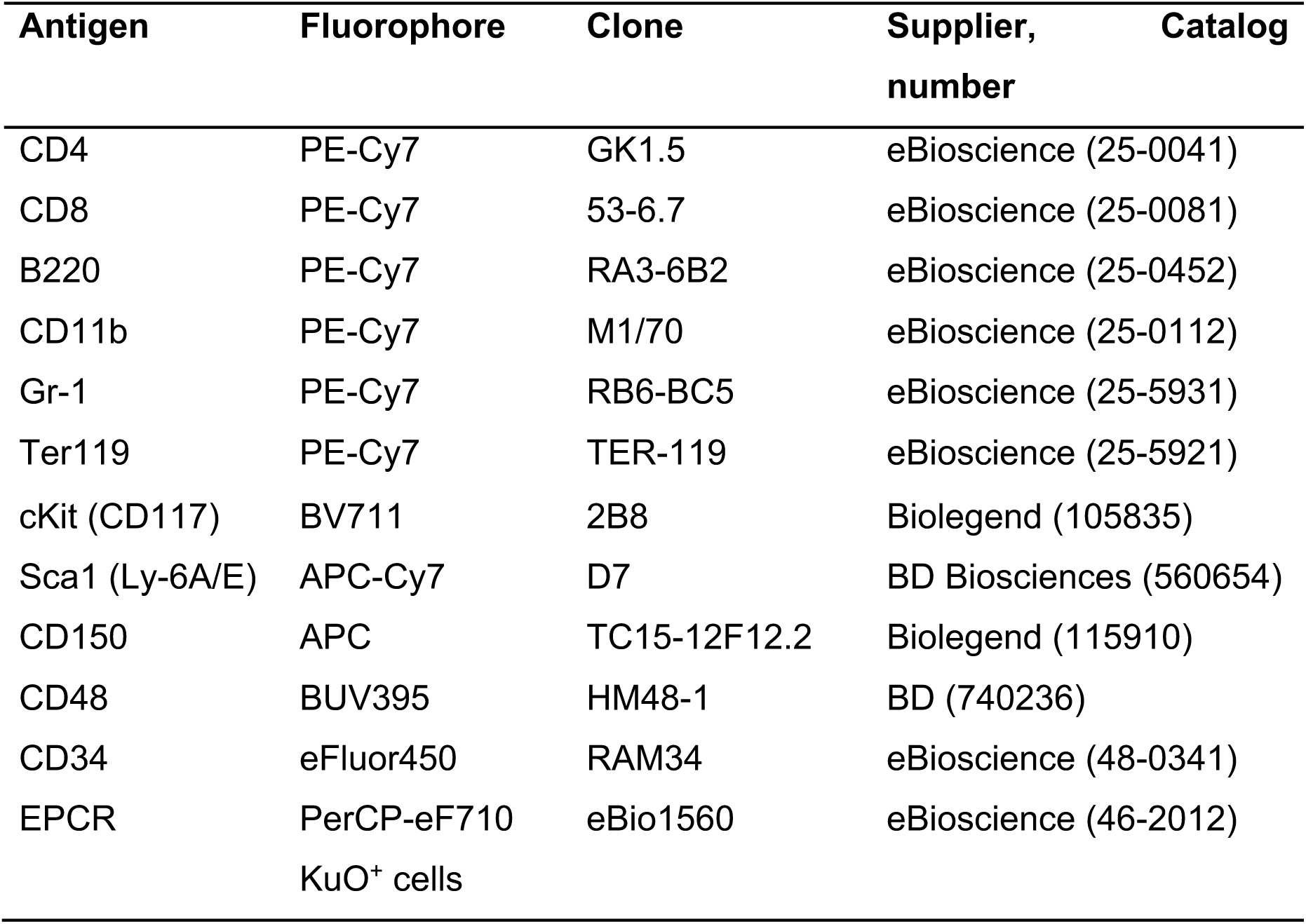
Antibody panel for the isolation of KuO^+^ EPCR^hi^ LT-HSCs.

### Antibody-based staining of hematopoietic cells

Peripheral blood, bone marrow, spleen, lymph nodes, liver, lung, thymus, colon and peritoneal cavity cell suspensions were stained using monoclonal antibodies recognizing cell-specific surface proteins. Cells were incubated with an antibody mix prepared in PBS + 2% FBS. For organ-derived hematopoietic staining, cell suspensions had a concentration of 1*10^5^ cells/µL antibody mix. For white blood cell staining, 50 µL peripheral blood was incubated with 100 µL antibody mix. Blood platelet and erythrocyte staining involved 3 µL peripheral blood and 27 µL antibody mix. Cells were incubated for 30 min at 4°C in the dark. All samples stained with antibodies against white blood cell epitopes were subjected to an erythrocyte lysis step using an ACK lysis buffer. Blood cells were incubated with ACK lysis buffer for 10 min, and remaining organ-derived hematopoietic cells were incubated with ACK lysis buffer for 2 min at room temperature. In case of the platelet and erythrocyte staining, this lysis step was not performed. After the lysis, cells were washed once with PBS + 2% FCS and resuspended in a final volume of PBS + 2% FCS. All samples were filtered prior to flow cytometry analysis.

### Flow cytometry analysis

Cells were analyzed by flow cytometry using a LSRFortessa or a LSRII cytometer (BD Biosciences), both equipped with 350 nm, 405 nm, 488 nm, 561 nm and 641 nm excitation lasers. Each antibody panel was manually compensated using OneComp eBeads (eBioscience) stained with single antibodies.

### Data pre-processing

Flow cytometry data were initially analyzed in FlowJo (v10.6.1, BD). Each defined cell population was divided into their parental congenic origin GFP+ CD45.1+ (donor), CD45.1/2+ (supportive bone marrow) and CD45.2+ (recipient) and the cell count, or frequency of parent (FoP) was imported into R (v4.1). For count data, percent relative donor engraftment (DE) per cell population was calculated as follows: DE = #donor / (#donor+#supportive+#recipient). For frequencies, the FoP of donor-derived cells corresponded to DE. To account for technical noise, the lower bound detection limit was adjusted for by setting the cell populations’ DE with less than 20 detected events to NA and the DE of less than 0.1% to 0%. Further, cell populations that did not reach the detection threshold in at least 10 analyzed samples were excluded from downstream analysis. For peripheral blood reconstitution analysis of the discovery cohort, mice were excluded if they experienced graft failure post-transplantation or did not reach an overall DE (i.e. donor chimerism) of greater than 0.1% at any time point. For final time point analysis of the discovery cohort, mice were excluded if they did not reach sustained DE of at least 0.1% in any PBMC sample at week 20 post-transplant.

### Dimensionality reduction and clustering

For each clonally-derived system, filtered DE levels of each organ-specific cell type were transformed into compositions. Prior to regression analysis, missing values were imputed by their mean. Dimensionality reduction was performed by principal component analysis (PCA) and the top 3 dimensions were chosen for hierarchical clustering on principal components (HCPC) using the FactoMineR (v2.6) package. For comparison of the generated clusters with previously defined HSC subtypes, hematopoietic systems were classified as described in Dykstra et al.^6^ and visualized using ggtern (v3.4.2).

### Relative repopulation capacity

The relative repopulation capacity of each hematopoietic system was calculated by dividing the overall peripheral blood chimerism levels (filtered DE levels of all blood cell types per system) from the secondary transplantation by its corresponding chimerism levels from the primary transplantation per week.

### Model fitting

Hematopoietic reconstitution kinetics were modeled by fitting the filtered DE levels of blood cells per hematopoietic system for each available time point using the ‘single humped function’ that is described as: x(t) = 0, if t < τ; x(t) = A * (t - τ) / (1 + ((t - τ) / θ) ^ n), if t >= τ, where τ is the delay, A the amplitude, θ the repression coefficient, and n the Hill coefficient. Parameter fitting was performed in Julia (v1.6) using the ModelFitter package (https://github.com/vkumpost/ModelFitter). Curve-specific characteristics (kinetic parameters) for each fitted curve were calculated as follows:

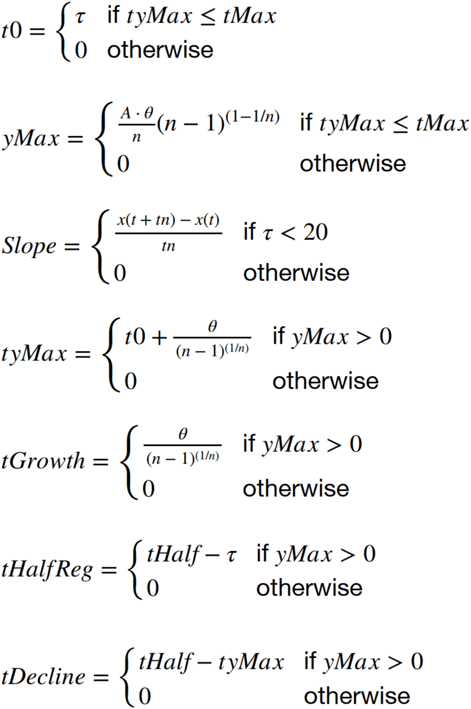

tHalf needed to be estimated using the Gauss-Newton method for non-linear systems. This was done by newtonsys(Ffun = x(t) – 0.5*yMax, x0 = tyMax+x0) from pracma (v2.3.8). The AUC for each fitted curve was calculated using the auc function from flux (v.0.3). Fits were excluded if the RMSE was > 0.08. All kinetic parameters except yMax, tyMax and AUC were set to NA if no decline was observed at the end of the study (tMax). Parameters yMax, tyMax and AUC were set to its value at tMax. All parameters were set to NA, if no chimerism was observed (t0 = tMax).

### Correlation analysis

Correlation analysis was performed using the rcorr() function from the Hmisc (v4.7-1) package. If not stated otherwise, Spearman rank correlation was used as a method. Polyclonal controls were excluded for these analyses. For visualization, either ComplexHeatmap (v2.10.0) or corrplot (v0.92) was used.

### HSPC transition

For each clonally-derived system, filtered DE levels of each HSPC were transformed into compositions. The compositions were ordered clockwise by their Pearson correlation distance to HSCs. The HSPC transition for each system was defined as the radian from HSC to median composition.

### Hierarchical clustering

Hematopoietic systems were clustered using hierarchical clustering on parameters t0 and AUC with Euclidean distance, ward.D2 as algorithm and k = 4 clusters (stats::hclust(), (v4.1.0)). Entanglement with clusters from PCA analysis was visualized and calculated using dendextend (v1.15.2). Kruskal Wallis test was used to assess significant differences between the kinetic parameters and the 3 groups for each blood cell type.

### Single-cell RNA sequencing and data preprocessing

For single-cell RNA sequencing, the Chromium Single Cell 3’ kit (v3.1) was used according to the manufacturer’s instructions. Libraries were sequenced on an Illumina HiSeq4000. FastQ files were processed and aligned using the Cell Ranger pipeline (v3.1) and the murine reference genome GRCm38 (mm10).

### Quality control and batch integration

Each individual sample was loaded into a SeuratObject (v4.0.4) using the Seurat framework (v4.1.0) for downstream analysis. cKit+ and total bone marrow (tBM) cells were filtered separately. cKit+ cells were kept if they had 700 – 6,000 features, 1,400 – 45,000 counts and less then 10% mitochondrial reads. tBM cells were retained if they had 300 – 5,500 features, 1,000 – 40,000 counts and less than 8% mitochondrial reads. The data were log-normalized, and the top 3000 variable features were scaled according to Seurat defaults. For data integration, LIGER was used via SeuratWrappers (v0.3.0) with default parameters, besides k = 50. Samples were treated as independent batches.

### Dimensionality reduction and clustering

The 50 factors generated from the data integration via LIGER were used for further dimensionality reduction into two-dimensional space using uniform manifold approximation and projection (UMAP), as well as for Louvain clustering with a final resolution of 0.9. Final annotation was performed based on known marker genes for each population.

### Differential abundance analysis

For differential abundance analysis, cell counts were transformed to compositions for each sample. Changes in abundance were assessed by calculating the log2-fold change difference between each clonally-derived cell type fraction and the corresponding polyclonal control fraction that was summarized as mean.

### Pseudotime analysis

Slingshot (v2.2.1) was used to calculate pseudotime trajectories for the progenitor compartment. The HSPC compartment was subset from the global dataset. The HSC cluster was chosen as the starting point and the distinct progenitors as endpoints. The UMAP was used as dimensionality input, on which the minimum spanning tree was calculated with default parameters. The curves were fitted using getCurves(extend = “n”, stretch =0).

### Modeling chimerism dynamics in mature blood populations

To investigate the differences in chimerism dynamics between fast and slow clonal systems in mature blood populations, an ordinary differential equation model was constructed. The model consists of three hierarchically arranged stem cell populations, subsequent progenitor populations and a final mature cell compartment. The number of populations downstream of stem cells was set to ten to account for progressive maturation of progenitor/precursor cells. Production of blood cells from the upstream compartments along the hematopoietic hierarchy is allowed by differentiation reactions. Chimerism dynamics were described by the following ordinary differential equation system:

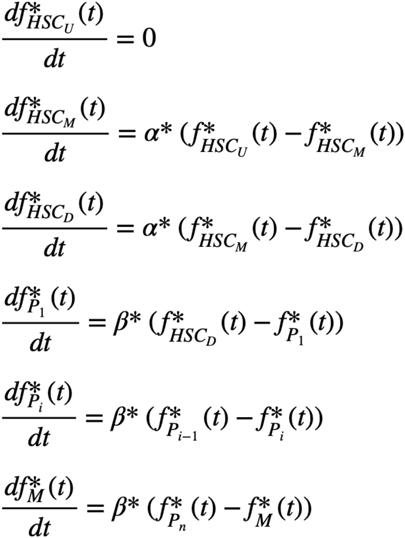

HSC_i_, P_i_ and M denote stem, progenitor and mature cell compartments, respectively, and f_Pi_(t) denotes chimerism in population P_i_. The model was separately fitted to average chimerism dynamics of seven hematopoietic lineages denoted by the asterisk: PLT, RBC, monocytes, granulocytes, B cells, CD4^+^ T cells and CD8^+^ T cells. For each lineage, average chimerism dynamics in slow and fast clusters were fitted simultaneously. The clusters were identified by hierarchical clustering of chimerism values in stem cells and mature lineages. Bayesian inference was employed to obtain posterior distributions of α and β using Turing.jl package (v0.24.0) in Julia (v1.8.5). For each differentiation step, involving a progenitor and a product population pair, α and β represent the product of the respective differentiation rate and compartment size ratio of the progenitor and product populations. Initial chimerism values were estimated for upstream and downstream stem cell populations in slow and fast clusters and set to zero for other populations.

### Mathematical modeling of the coefficient of variation in linear and feedback compartment models

To address our observation that cell counts in mature blood populations display significantly lower coefficients of variation than LT-HSCs, we simulated two compartment models. Both models consist of three populations: stem cells (S), progenitors (P) and mature cells (M). Stem cells proliferate with rate λ_S_ and differentiate into progenitor cells with rate δ_S_. Progenitor cells, in turn, proliferate with rate λ_P_ and differentiate into mature cells with rate δ_P_. Mature cells undergo cell death with rate δ_M_. In the linear model, all proliferation and differentiation fluxes are proportional to the respective population sizes. In the feedback model, progenitor cell proliferation is governed by negative feedback and is implemented using a carrying capacity for the progenitor population, P_C_; this ensures stable regulation of mature cell numbers The dynamics of the linear model are described by the following ordinary differential equation system:

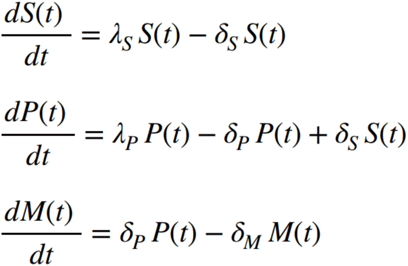

Similarly, the feedback model is described by the following nonlinear ordinary differential equation system.

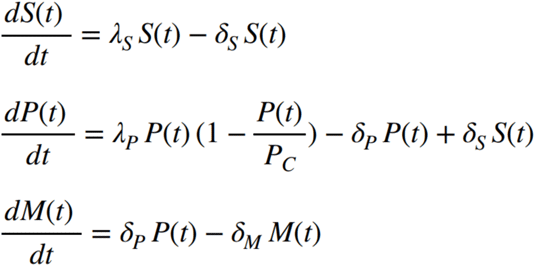

Simulations for both models were initiated with hundred cells in the stem cell compartment (S_0_ = 100, P_0_ = 0, M_0_ = 0) and propagated up to 300 days. Proliferation and differentiation rates of the linear compartment model were set to the following values: λ_S_ = 0.1, δ_S_ = 0.1, λ_P_ = 2.0, δ_P_ = 2.02, δ_M_ = 0.1. For the feedback model the following rates were used: λ_S_= 0.1, δ_S_ = 0.1, λ_P_ = 2.1, δ_P_ = 2.02, P_C_=10500, δ_M_ = 0.1. Coefficients of variation for individual compartments were computed from hundred independent simulations and normalized to the stem cell compartment. Simulations were performed with CoRC (v0.11.0, COPASI v4.34)^39,40^ in R (v3.6.1).

### Data visualization and statistical analysis

If not specifically stated otherwise, significance was tested using paired samples Wilcoxon test. For multiple comparisons, p-values were adjusted according to Benjamini & Hochberg. Plots were generated using ggplot2 (v3.4.2) or FlowJo (v10.6.1).

## Supplementary Figures

**Supplementary Fig. 1:**
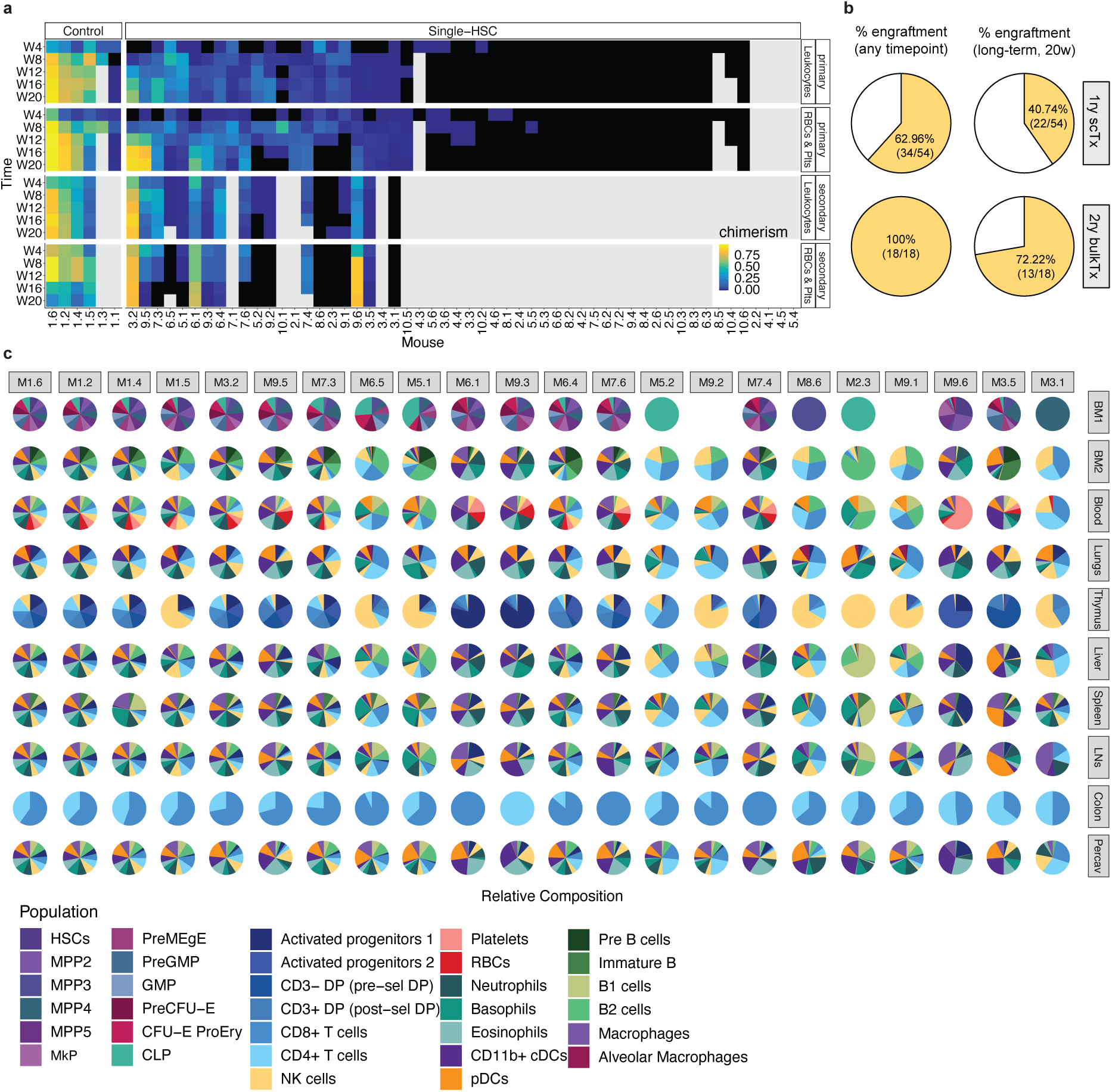
Overview of single-HSC transplantation experiment. **a)** Heatmap of donor chimerism over time split between overall leukocyte, and erythrocyte and platelet chimerism of transplanted mice within primary and secondary transplantation ordered by their overall chimerism. Black color indicates chimerism <0.1%, gray color indicates missing value. **b)** Pie charts displaying the percentage of clones with successful engraftment (>0.1% chimerism in peripheral blood) at any time point post-transplantation (left), and with positive long-term chimerism (at 20 weeks post-transplantation) (right), for primary (top) and secondary (bottom) transplants. 1ry scTx: primary single-cell transplantation; 2ry bulkTx: secondary bulk bone marrow transplantation. **c)** Clonal hematopoietic compositions 20 weeks post-transplant split per organ and cell types are highlighted by color. BM1 and BM2 correspond to two different antibody stainings of the bone marrow. Abbreviations: HSC: hematopoietic stem cell; RBCs: red blood cells; Plts: platelets; W: week; BM: bone marrow; ScTx: single-cell transplantation; LNs: lymph nodes, MPP: multipotent progenitor; MkP: megakaryocyte progenitor; PreMegE: pre-megakaryocyte-erythrocyte; GMP: granulocyte-monocyte progenitor; PreCFU-E: pre-colony-forming-unit-erythrocyte; CFU-E-ProEry: colony-forming-unit-erythrocyte-proerythroblast; CLP: common lymphoid progenitor; DP: double positive; cDC: conventional dendritic cell; pDC: plasmacytoid dendritic cell; NK: natural killer cell.

**Supplementary Fig. 2:**
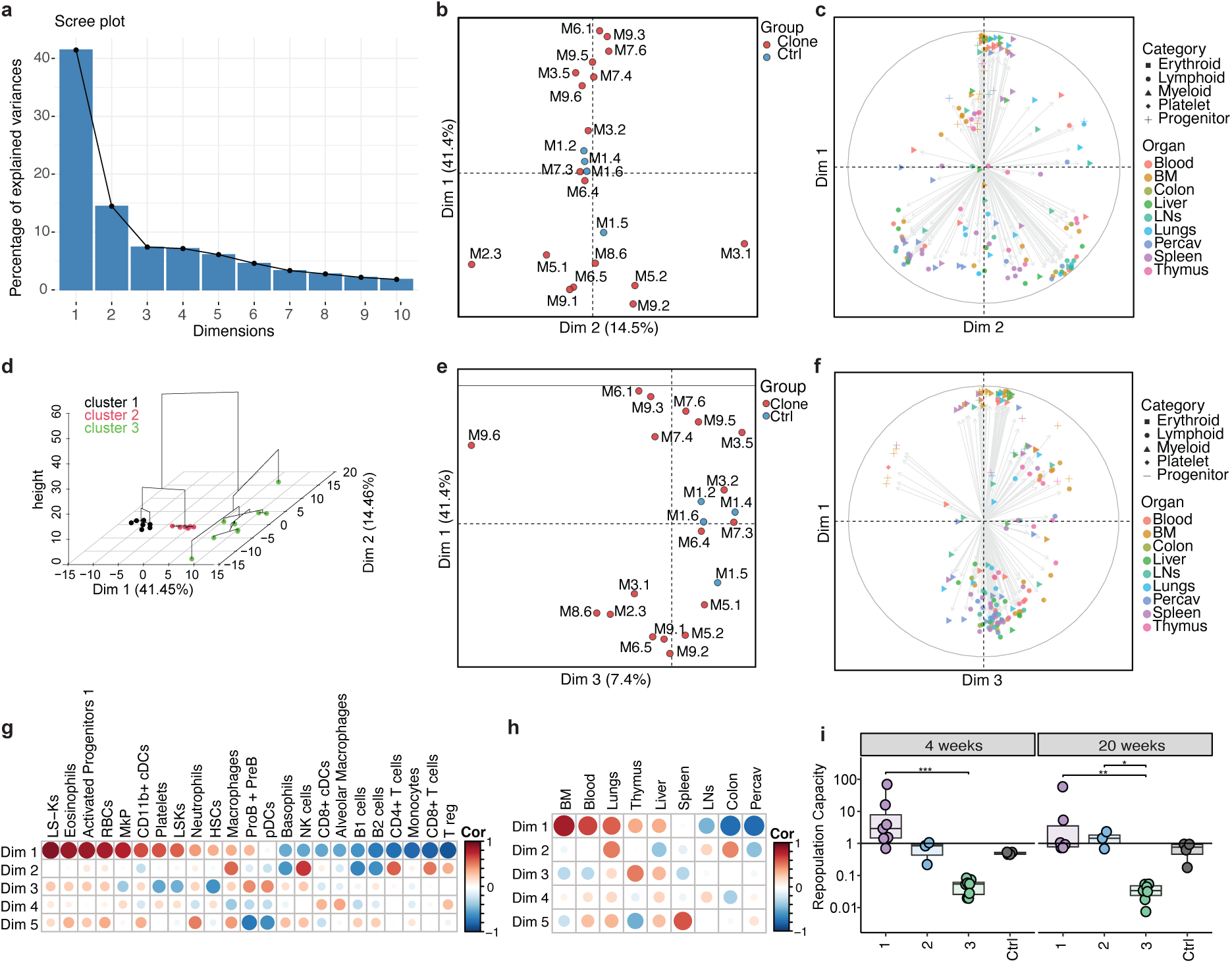
Principal component analysis of clonally-derived hematopoietic systems. **a)** Scree plot showing the percentage of explained variance by each principal component. **b)** First and second principal component projections. Individuals are labeled by experiment ID and colored by group (single HSC-derived: red, polyclonal control: blue). **c)** Variable contribution map explaining the distribution of individuals between the first two dimensions based on organ tropism and lineage contribution. **d)** Hierarchical clustering on principal components based on the first three PCs. **e)** First and third principal component projections, see (b). **f)** Variable contribution map of first and third principal component projections, see (c). **g-h**) Spearman correlation coefficients between active and supplementary variables and dimensions, highlighting associations between principal components and cell types (g) as well as organs (h). **i)** Relative repopulation capacity (defined by the ratio of chimerism in secondary and primary transplantations) across the three clusters defined in Fig. 1, and polyclonal controls based. If not stated otherwise, significant differences between groups were tested globally by Kruskal Wallis test and post hoc by two-sided Wilcoxon rank-sum test. For multiple comparisons, p values were corrected according to Benjamini-Hochberg. Significance is indicated by: * for p < 0.05, ** for p < 0.01, *** for p < 0.001. The standard deviation is indicated by error bars. Box plots: center line, median; box limits, first and third quartile; whiskers, smallest/largest value no further than 1.5*IQR from corresponding hinge. Abbreviations: Ctrl: control; HSPC: hematopoietic stem and progenitor cell; cDC: conventional dendritic cell; pDC, plasmacytoid dendritic cell; NK cell: natural killer cell; RBC: red blood cell; BM: bone marrow; LN: lymph node; PerCav: peritoneal cavity; ctrl: control; dim: dimension; LS-K: Lineage-Sca1-cKit+; MkP: megakaryocyte progenitor; LSK: Lineage-Sca1+cKit+.

**Supplementary Fig. 3:**
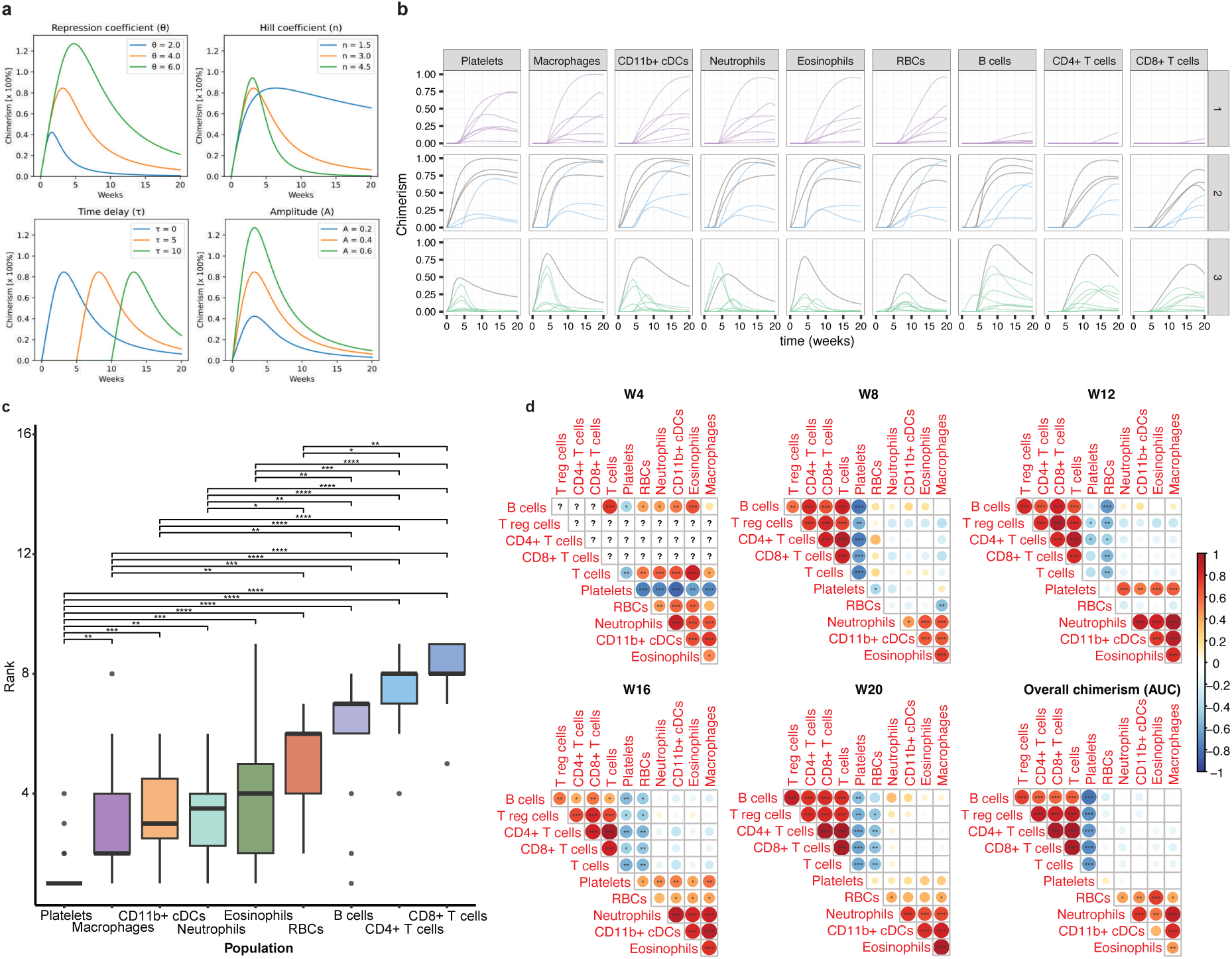
Kinetic curve fitting parameters. **a)** Exemplary behavior of single humped function fits with variation in its four coefficients. Repression coefficient and hill coefficient describe how stretched, or compressed the curve declines, the time delay marks the initial time point of growth and the amplitude controls the initial growth of the curve. **b)** Reconstitution kinetics of peripheral blood cells from transplanted HSCs separated by hierarchical clusters and cell type. Each coloured line corresponds to a single clonally-derived system. Polyclonal controls are colored in gray. **c)** Blood cells ordered by their ranked engraftment delay t0 within clonally-derived hematopoietic systems. Significant differences between cell types were tested globally by Kruskal Wallis test and post hoc by Dunn’s test. For multiple comparisons, p values were corrected according to Benjamini-Hochberg. **d)** Spearman correlation matrices of clonal cell type compositions within the blood. Correlations of single HSC-derived blood cell compositions were calculated for each measured time point, or using the overall chimerism values derived from AUC of each fitted curve, respectively. Cell types are ordered by angular order of eigenvectors from the correlation of AUC compositions. Significance is indicated by: * for p < 0.05, ** for p < 0.01, *** for p < 0.001, **** for p < 0.0001. Box plots: center line, median; box limits, first and third quartile; whiskers, smallest/largest value no further than 1.5*IQR from corresponding hinge. Dots indicate outliers. Abbreviations: cDC: conventional dendritic cell; RBC: red blood cell; W: week; AUC: area under the curve.

**Supplementary Fig. 4:**
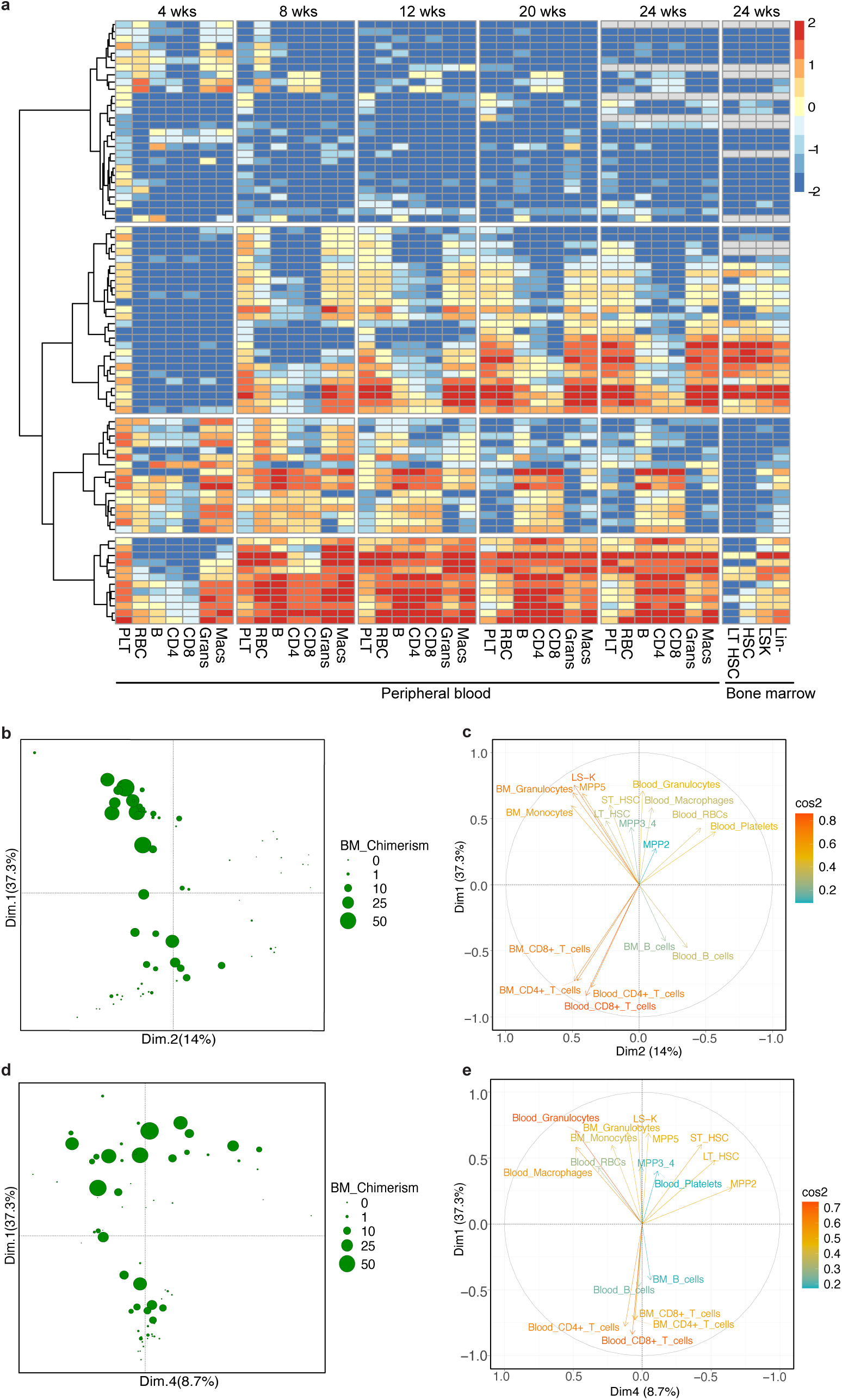
Validation cohort confirms reconstitution kinetics as a central feature associated with HSC potency and lineage biases. **a)** Heatmap displaying the percentage of chimerism in peripheral blood at different timepoints post-transplantation and in the bone marrow at the endpoint (24 weeks). Chimerism values are presented as log₁₀-transformed percentages. Each row represents a single HSC transplantation that displayed positive chimerism (>0.1% in peripheral blood) at any time point. Agglomerative clustering, named AGNES (AGglomerative NESting) was used. **b)** Principal component analysis (PCA) considering the cellular composition of all HSC-derived cell types at the endpoint of the primary transplant (week 24). The first two components are displayed and overall chimerism is highlighted by dot size. **c)** Variable contribution map of (b) highlighting the loadings by differentiation status and lineage. **d)** First and fourth principal component projections of PCA from (b). Overall chimerism is highlighted by dot size. **e)** Variable contribution map of (b) highlighting the loadings by differentiation status and lineage from (d). Abbreviations: wks: weeks; PLT: platelet; RBC: red blood cell; B: B cell; CD4: CD4+ T cell; CD8: CD8+ T cell; Grans: granulocytes; Macs: macrophages; LT-HSC: long-term hematopoietic stem cell; LSK: Lineage-Sca1+cKit+; Lin-: lineage negative; BM: bone marrow; Dim: dimension; MPP: multipotent progenitor; ST-HSC: short-term hematopoietic stem cell.

**Supplementary Fig. 5:**
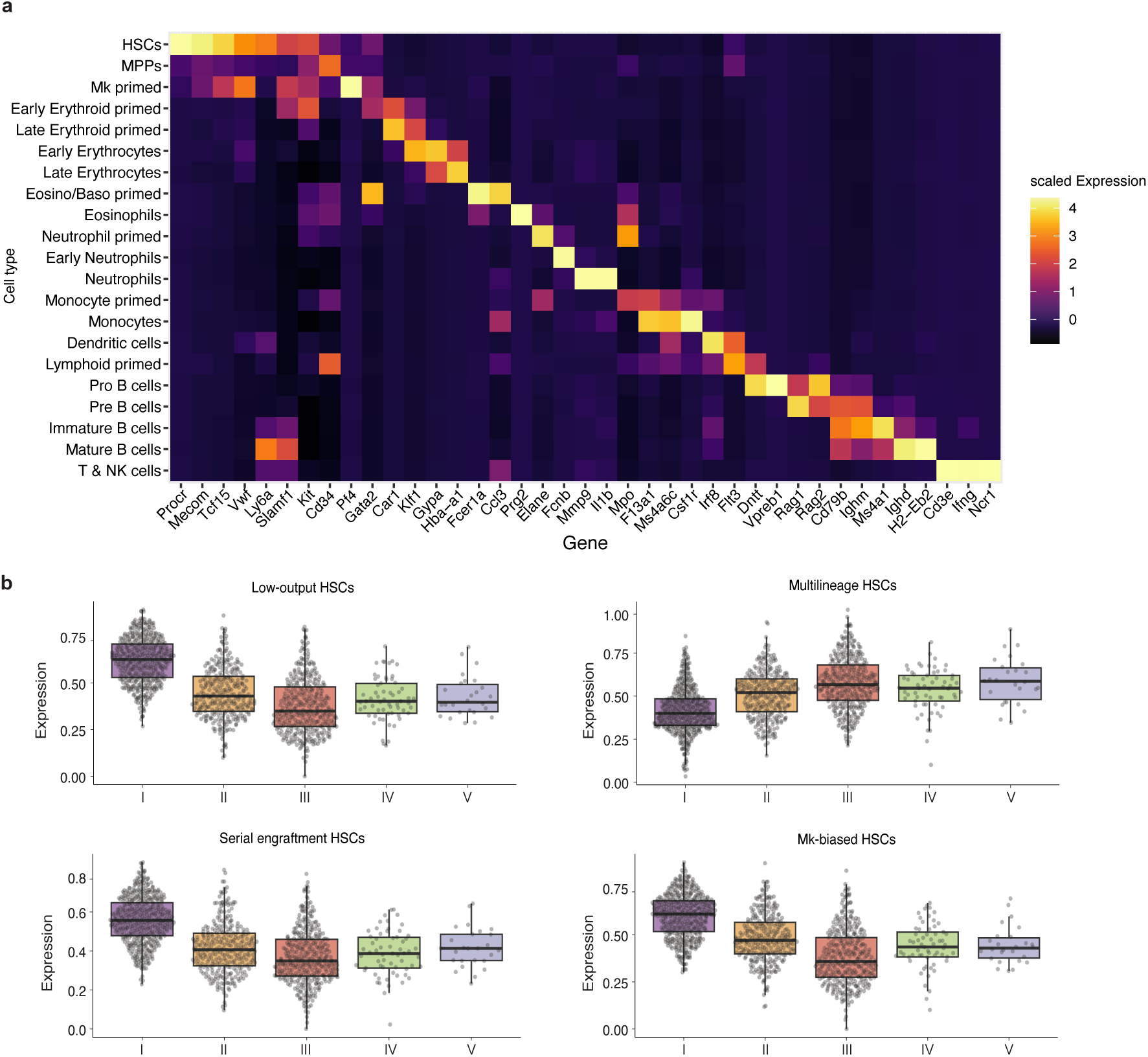
Single-cell transcriptomics analysis of clonally-derived hematopoietic systems. a) Gene expression heatmap illustrating major marker genes for each progenitor and cell type cluster. The color scale highlights the scaled average expression per gene in each cluster. b) Boxplots highlighting module scores of gene sets characteristic for low-output HSCs, multilineage HSCs, serial engrafting HSCs and Mk-biased HSCs in each HSC per clonally-derived system ordered by increasing blood cell repopulation and decrease in self-renewal. Gene sets are derived from^8^. Box plots: center line, median; box limits, first and third quartile; whiskers, smallest/largest value no further than 1.5*IQR from corresponding hinge. Abbreviations: HSC: hematopoietic stem cell; MPP: multipotent progenitor; Mk: megakaryocyte; Eosino: eosinophil; Baso: basophil; NK: natural killer cell.

**Supplementary Fig. 6:**
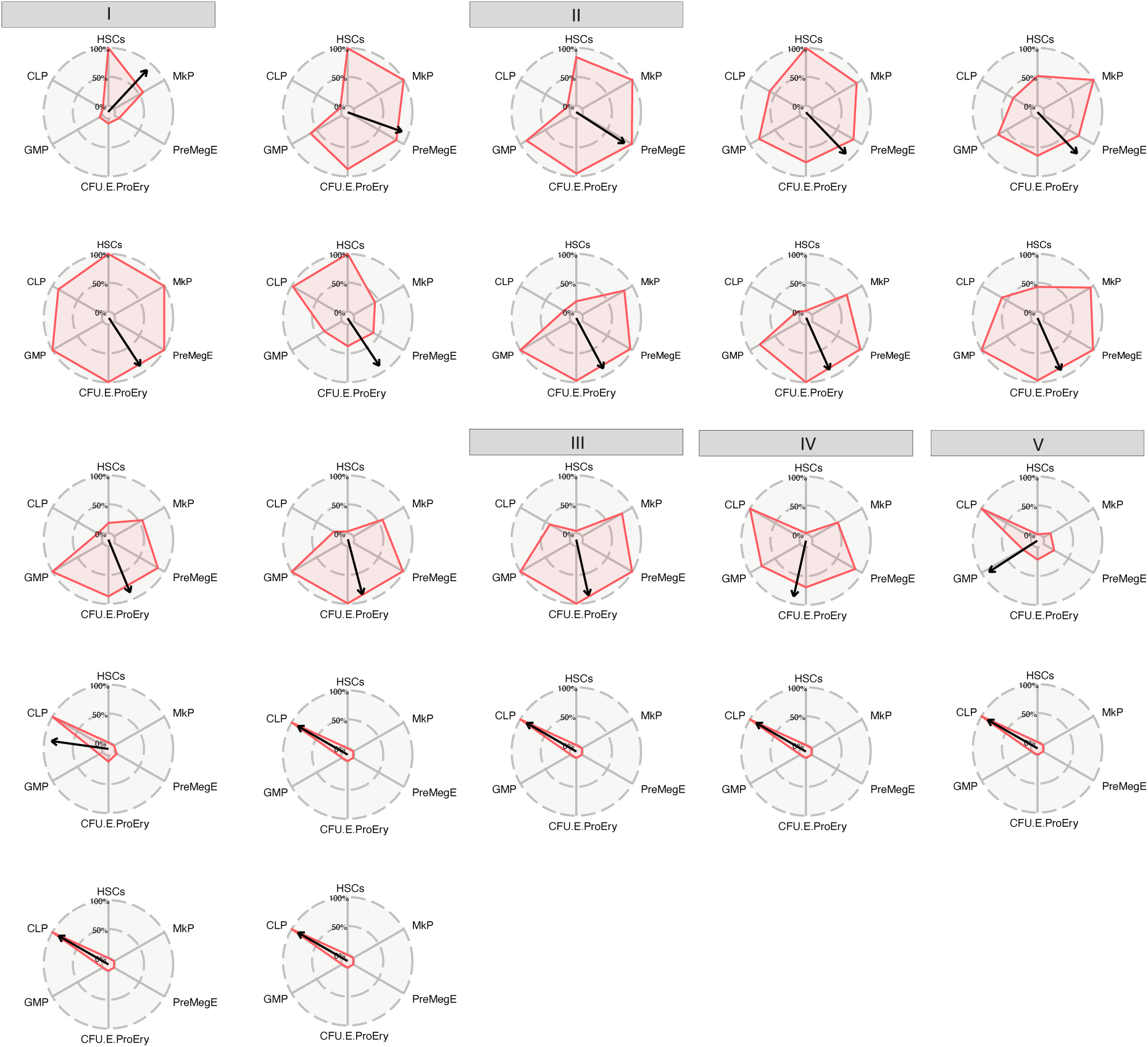
HSPC transition clocks for clonally-derived hematopoietic systems. Composition of HSPC compartments of clonally-derived systems from Fig. 1 that displayed sustained engraftment. Progenitors are ordered by Pearson correlation distance from HSCs based on clonal compositions of the progenitor compartments 20 weeks post-transplant (see illustration Fig. 3f). The arrow indicates the mean composition of the respective clonally-derived HSPC compartment and illustrates the current state of “HSPC transition”. Clonally-derived systems marked as I, II, III, IV and V correspond to the exemplary plots shown in Fig. 3f. Abbreviations: HSC: hematopoietic stem cell; MkP: megakaryocyte progenitor; PreMegE: pre-megakaryocyte-erythrocyte; CFU-E-ProEry: colony-forming-unit-erythrocyte-proerythroblast; GMP: granulocyte-monocyte progenitor; CLP: common lymphoid progenitor.

**Supplementary Fig. 7:**
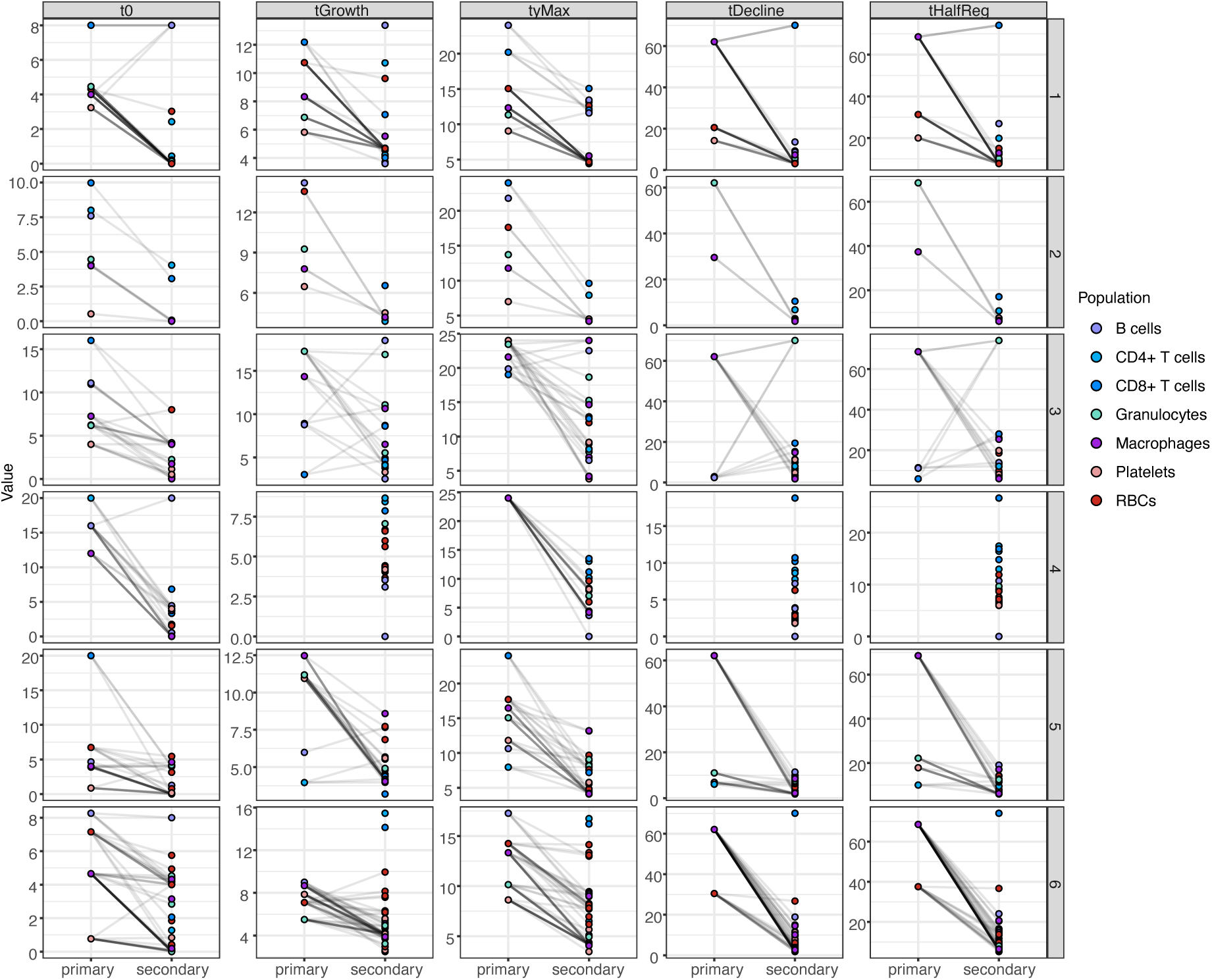
HSCs transition from slow to fast reconstitution kinetics upon secondary transplantation. Paired dot-plots of time-dependent kinetic parameters highlighting changes in blood cell replenishment between transplanted parent and daughter HSCs per clone. Cell types are highlighted by color. Each row represents a paired analysis between parent (primary) and corresponding daughter (secondary) HSCs. Abbreviations: RBC: red blood cell.

**Supplementary Fig. 8:**
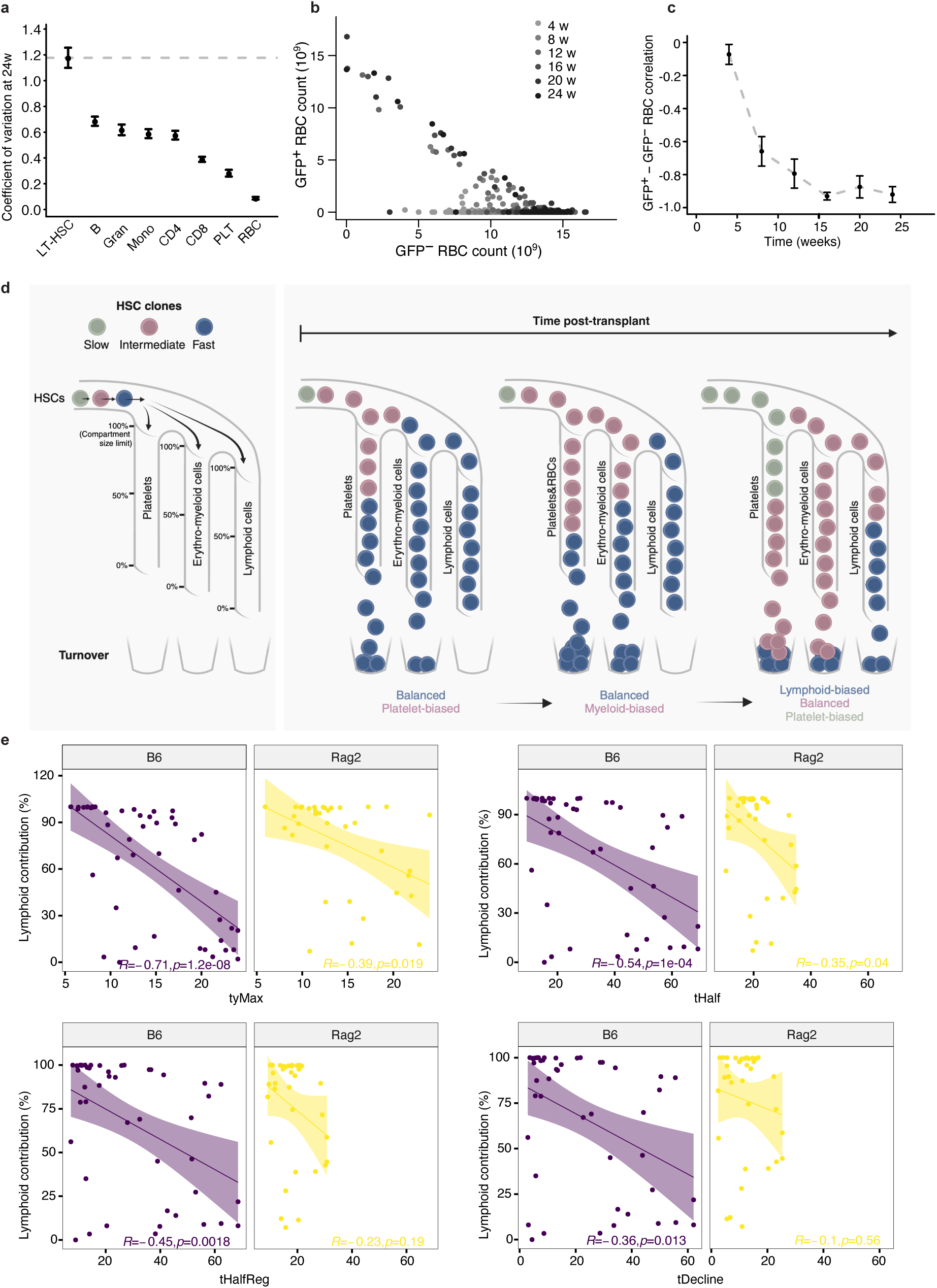
Cell-extrinsic mechanisms modulate HSC lineage biases. **a)** Coefficient of variation of bone marrow LT-HSCs and peripheral blood B cells (B), granulocytes (Gran), monocytes (Mono), CD4+ T cells (CD4), CD8+ T cells (CD8), platelets (PLT) and red blood cells (RBC) at 24 weeks post-transplantation. **b)** Correlation analysis between the absolute counts of single HSC-derived GFP+ and non-clonal GFP-RBCs measured in the peripheral blood at 4, 8, 12, 16, 20 and 24 weeks post-transplantation. **c)** Correlation coefficient extracted from (b) at different time points post-transplantation. **d)** Schematic representation of the proposed relationship between HSC clonal reconstitution kinetics, mature lineage compartment size limitations and lineage skewing. Three HSC clones with differing kinetics are depicted. The fast clone rapidly populates all mature blood lineages up to their compartment size limits. After the HSPC compartment of the fast clone exhausts, the mature cell progeny decline according to the rate of turnover of each lineage, meaning that the intermediate kinetics HSC clone first has space to populate the platelet and RBC compartment, then the myeloid compartment and finally the lymphoid compartment. Eventually the HSPC compartment of the intermediate clone exhausts, resulting in its progeny being sequentially replaced by those of the slow kinetics HSC clone. The resulting apparent lineage biases are indicated at three different time points post-transplantation. **e)** Spearman correlation between the average kinetics parameters tyMax, tHalf, tHalfReg and tDecline of a single HSC and its percentage of lymphoid contribution in peripheral blood at 24 weeks post-transplantation, in B6 versus Rag2^-/-^ hosts. Spearman’s Rho and significance are indicated. Abbreviations: LT-HSC: long-term hematopoietic stem cell; B: B cells; Gran: granulocyte; Mono: monocyte; CD4: CD4+ T cell; CD8: CD8+ T cell; PLT: platelet; RBC: red blood cell; w: week; B6: C57BL/6J mouse model; Rag2: homozygous knock-out of the Recombination Activating Gene 2.

**Supplementary Fig. 9:**
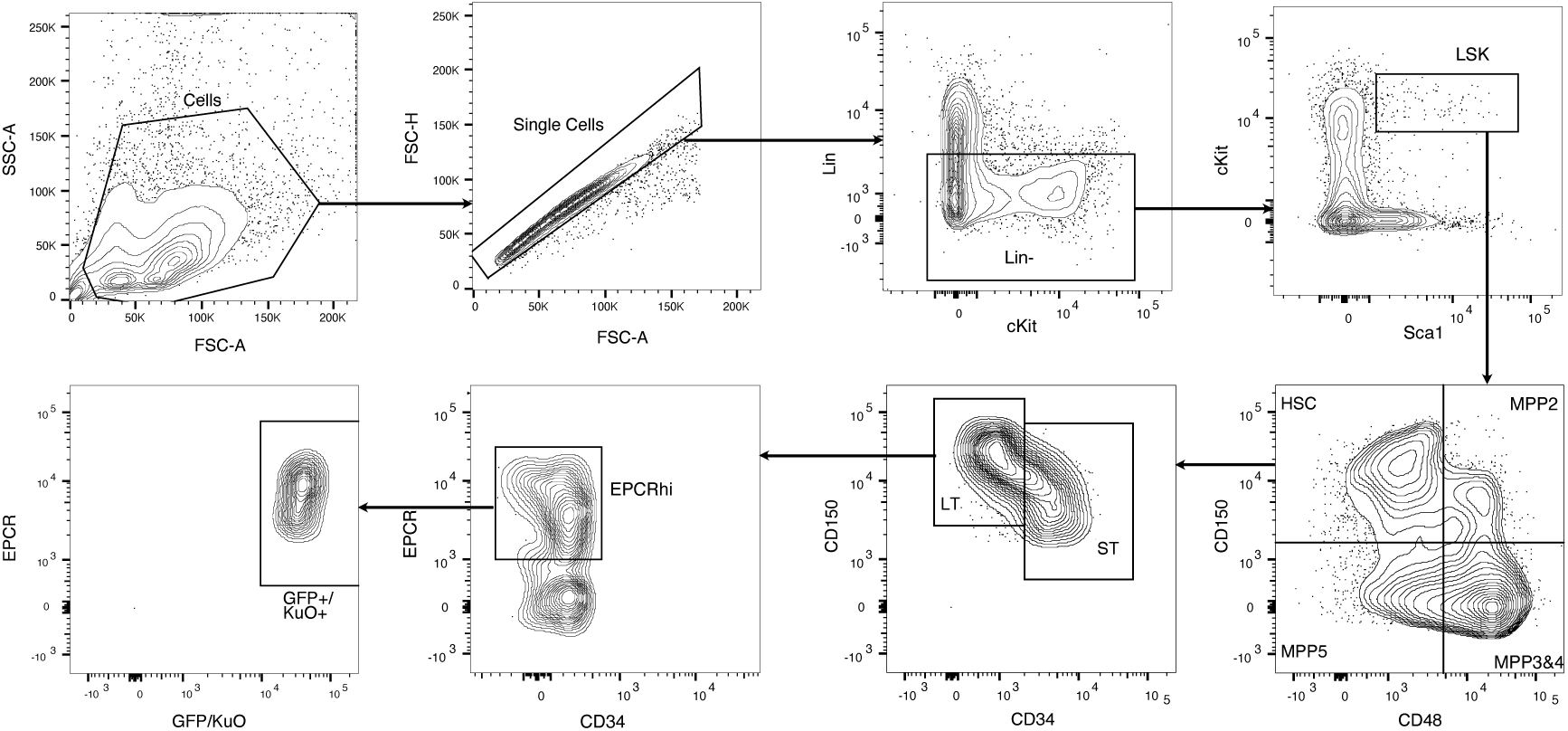
Gating strategy to sort donor EPCRhi LT-HSCs for transplantations studies.

